# Acid Stress-induced plasma membrane ceramide arises from lysosome-plasma membrane fusion

**DOI:** 10.64898/2026.09.21.753246

**Authors:** Abhay Kanodia, Anne Ostermeyer-Fay, Maria Hernandez-Corbacho, Ryan Chacon, Mehdi Damaghi, Yusuf A. Hannun, Daniel Canals

## Abstract

Plasma membrane ceramide (PMCer) is a recognized cellular signal that regulates stress responses, such as chemotherapy and membrane damage, triggering apoptosis and plasma membrane repair. Acid sphingomyelinase (ASM) has been proposed to generate PMCer by directly hydrolyzing plasma membrane sphingomyelin into ceramide. However, the enzymatic requirements of ASM are difficult to reconcile with the extracellular environment on the outer leaflet of the plasma membrane. Here, we identify several cellular stress inducers, with acidic stress yielding a stronger response in PMCer generation. Using this inducer, we defined a lysosome-to-plasma membrane pathway for stress-induced PMCer generation. Acidic stress increased PMCer in an ASM-dependent manner, whereas ASM loss or catalytic inactivation abolished this response. Although secreted ASM was enzymatically active when tested in vitro, neither secreted nor exogenous recombinant ASM generated PMCer. Instead, acidic stress induced lysosomal exocytosis, while disruption of lysosomal fusion reduced PMCer accumulation. Modulating lysosomal ceramide translated into PMCer, showing that lysosomal ceramide is directly transferred into the plasma membrane. These findings redefine stress-related PMCer signaling through lysosomal ceramide exocytosis.

## INTRODUCTION

Research over the past four decades has demonstrated that sphingolipids—particularly ceramide and sphingosine 1-phosphate—play critical roles in a wide range of biological processes. Ceramide occupies a central position in sphingolipid metabolism and has been implicated in apoptosis, autophagy, senescence, and cell cycle arrest, thereby contributing to the pathogenesis of diseases such as cardiovascular disorders, cancer, and neurological conditions [1].

Notably, studies of ceramide biology have revealed distinct functions depending on cellular context. These observations led to the formulation of the “Many Ceramides” hypothesis, articulated by Hannun and Obeid in 2011 [2]. The hypothesis reflects a growing consensus that ceramide should not be considered a single entity but rather a family of structurally diverse species, compartmentalized within cells, and metabolized by distinct enzymatic pathways. This spatial and metabolic organization is thought to dictate the specific biological functions of individual ceramide pools.

A few enzymes have been studied in the context of bioactive plasma membrane ceramide (PMCer) generation. Neutral sphingomyelinase 2 (nSMase2) has been shown to regulate basal levels of PMCer as well as respond to physiological stimulation (such as cell adhesion and cell migration). On the other hand, acid sphingomyelinase (ASM) has been associated primarily with stress-related stimuli, including radiation, viral or bacterial infection, exposure to Fas ligand (FasL), and plasma membrane repair [3–6]. Other enzymes have been shown to act on the plasma membrane (SMS2 and GBA2), but their contribution to PMCer or ceramide-related biology has not been studied in detail [7].

Across stress conditions, a prevailing model proposes that intracellular ASM translocates to the plasma membrane, where it generates ceramide, triggering downstream signaling events and biological outcomes, including cell death, facilitation of infection, and plasma membrane repair [8, 9]. However, this prevailing model of ASM-dependent PMCer generation is difficult to reconcile with several enzymatic and biochemical constraints. These include uncertainty regarding (i) how ASM traffics to and is retained at the plasma membrane, (ii) how secreted ASM remains catalytically active towards plasma membrane sphingomyelin at near-neutral pH, and (iii) how it functions in the absence of key lysosomal cofactors. These cofactors include anionic lipids required for membrane binding, saposin C, which facilitates sphingomyelin hydrolysis under suboptimal conditions, and zinc ions required to activate the secreted enzyme.

One major limitation in studying PMCer is the lack of tools to probe and measure it. Previous work has reported using ceramide-specific antibodies, which do not enable mass quantification and are limited to qualitative ceramide detection. Moreover, many of these anti-ceramide antibodies have been shown to bind other lipids as well [10]. Recently, our laboratory developed a sensitive method to directly quantify PMCer [11]. Using this approach, the present work tracked the source of PMCer in response to distinct reported cellular stresses. Acidic stress generated the largest amount of PMCer and was chosen to study its generation mechanism. The results attribute this ceramide pool to ASM, confirming previous reports. However, our data contradict ASM acting on the plasma membrane, and we propose that ASM-derived ceramide located in the lysosome reaches the plasma membrane through lysosomal exocytosis.

We propose a novel cellular pathway that explains how ASM can drive the acute generation of bioactive ceramide at the plasma membrane during stress signaling without acting directly at the plasma membrane, where conditions are unfavorable for its enzymatic activity.

## Materials and Methods

### Cell lines

MCF-7 (HTB-22, an epithelial cell line from breast adenocarcinoma) was purchased from ATCC. MCF-7 was maintained in Roswell Park Memorial Institute (RPMI) 1640 medium from Thermo Fisher. Growth media were supplemented with 12.5 mM HEPES and PIPES, and 1 N NaOH was added to adjust the pH to 6.4 [12]. The cells were maintained at 37°C in a 5% CO_2_-supplied incubator. Mycoplasma testing was performed monthly.

### Plasmids

All the plasmids, Niemann-Pick mutants (R498L and L304P) in ASM, and the catalytically inactive ACD mutant (C143A) sequences were cloned into pTwist Lenti SFFV Puro WPRE at Twist Biosciences. WT ASM and the mutant S508A were previously cloned into pLenti 6.3/V5 DEST [13].

### Sphingolipid quantification

MCF-7 cells were seeded in a 60 mm dish at 500K cells/dish in either neutral (pH 7.4) or acidic (pH 6.4) media. The medium was changed every 24 hours, and 24 hours later the cells were harvested by adding 2 ml of organic solvent (Ethyl acetate: isopropanol, 2:3) and 50 pmol of internal standards (non-natural d17-sphingolipids). The cells were scraped in the organic solvent and added to the glass tubes. Samples were centrifuged at 3000 x g for 5 minutes. The lipid extracts were transferred to a new tube and evaporated under nitrogen gas at a 42°C water bath. The dried film of lipids was brought to solution with 150 μl methanol, transferred to mass spectrometry vials, and stored at -20°C until injection into the LC-MS/MS analysis.

### PMCer quantification

PMCer was measured as previously described [11]. Briefly, 500k cells were seeded in 60 mm dishes and allowed to grow in neutral or acidic media for 24 hours. The cells were then washed twice in serum-free media and fixed with 8% paraformaldehyde (PFA) for 20 mins at room temperature. The dishes were divided equally, and half were treated with pCDase while the other half were untreated and incubated for 1 hour at 37**°**C in 5% CO_2_. The cells were then washed twice in serum-free media and collected in organic solvent (as described above). Sphingosine levels were measured using LC-MS/MS. PMCer was measured by subtracting pmol of sphingosine in the presence or absence of pCDase treatment. The PMCer was then normalized to the protein amount measured by BCA analysis.

### Immunofluorescence

Cells were plated in RPMI/10% FBS on poly-D-lysine-coated dishes (MatTek, Ashland, MA) at 10^5^ cells/ dish. After overnight in serum free media, cells were fixed (4% formaldehyde in PBS for 20 min), permeabilized (0.1% Triton X-100 in PBS, 10 min), blocked (3% BSA in PBS, 1 hour), and stained for 1 h at room temperature with either anti-V5 antibody (1: 500; Invitrogen; Cat # 46- 1175); anti-LAMP2 (1: 50; Santa Cruz Biotechnology, sc-18822), or anti-ASM (1:20; R and Biosystems; AF5348). After three washes with PBS, cells were incubated with goat anti-mouse 488 nm antibody (dilution 1:500) and goat anti-rabbit 500 nm antibody (dilution 1:500) in PBS for 1 h at room temperature. Following three washes with PBS, MatTek dishes were stored at 4 °C prior to image collection. All confocal images were taken with a laser scanning confocal microscope, Leica TCS SP8, with an HC PL APO 63×/1.40 oil immersion objective (Morrisville, NC).

### siRNA screening

The following siRNAs were used: human *SMPD1* siRNA (s13167), human SMPD3 siRNA (s30927) human GBA1 siRNA (s501316), human GBA2 siRNA (s33634), human SMS2 siRNA (Qiagen; 102720), human *ASAH1* siRNA (s496), *PTP4A3* siRNA (s104958), *MCOLN1* siRNA (s32877), *MCOLN3* siRNA (ID: s30633), and All Star. siRNA was used at 40 nM for transfection of cell lines, performed using Lipofectamine RNAiMAX Transfection Reagent (Thermo Fisher) via reverse transfection.

### Western blot

Cells were scraped in 250 μl of 2X Laemmli buffer (Bio-Rad) supplemented with 2- mercaptoethanol. Laemmli buffer (6X) (Thermo Fisher Scientific) was added to the conditioned media. The lysates or the conditioned media in Laemmli buffer were sonicated using a probe sonicator (2 pulses, 10 seconds each) and boiled at 100°C for 10 min and loaded on the gel (Bio-Rad 4–20% polyacrylamide Tris-Glycine gel). After proteins were separated on the gel based on size, they were transferred to a nitrocellulose membrane (1h, 4 °C). The membrane was blocked with 5% fat-free milk in Tris-phosphate buffer saline (TPBS) for 1 hour and then incubated with the primary antibody in 5% fatty-acid-free bovine serum albumin + 0.02% azide in TPBS overnight at 4°C. The antibodies used were anti-V5 (1:2000 Life Technologies, Cat # 46-1175), anti-ASM (1µg/ml, R&D Biosystems, Cat #AF5348), anti-ACD (1: 200; BD; cat # 612302). anti β-actin (1:10,000; Life technologies; cat. #A5441) was used as a loading control. Membranes were washed three times with TPBS and incubated for 1 hour with appropriate HRP conjugated secondary antibody either mouse (1:5000), Rabbit (1:5000), Goat (1:5000) in 5% Milk/TPBS. Western blots were developed with ECL and exposed on X-ray film.

### qRT-PCR

The RNA was purified using Invitrogen Pure Link RNA miniprep kit, RNA cat. #12183018A, and the cDNA was prepared using Invitrogen SuperScript III First-Strand Synthesis SuperMix for qRT-PCR kit, cat#11752050 following manufacturer’s instructions. TaqMan probes for human species were used for qRT-PCR: *ASAH1*, Hs00602774_m1; *GBA1*, Hs00164683_m1; *GBA2*, Mm00554547_m1; *SGMS2*, Hs00380453_m1; *SMPD1*, Hs03679347_g1; *SMPD3*, Hs00920354_m1. All gene expression was normalized using beta-actin (*ACTB*), Hs99999903_m1.

### ASM in vitro activity assay

The ASM activity was adapted from Jenkins et al. [14]. For lysosomal sphingomyelinase (L-ASM) activity, cells were harvested in L-SMase lysis buffer (0.2% Triton X-100, 50 mM Tris-HCl, pH 7.4. Brief sonication was carried out (1–2 pulses, 10 s), and cellular debris and unbroken cells were pelleted by centrifugation at 1,000 g for 5 min at 4 °C. 100 μl of clarified lysate was added to 100 μl of reaction mixture. The reaction mixture contained 100 μM sphingomyelin (d17:1/18:0) (Avanti, Alabaster, AL), in the assay buffer (0.2% Triton X-100 in sodium acetate buffer 250 mM, 1.0 mM EDTA, pH 5.0). Secreted ASMase (S-ASM) activity was performed with the indicated volume of conditioned medium in a final volume of 100 μl and assayed with a similar reaction buffer. ZnCl_2_ (0.1 mM) was substituted for 1 mM EDTA. For both S-ASM and L-ASM assays, the reaction was carried out for 10 min at 37 °C. Lipids were extracted by adding 1.8 ml of PBS and 2 ml of 15: 85 Isopropanol: Ethyl acetate to the reaction mix, followed by centrifugation at 3,000 x g for 5 min at room temperature to separate phases. The upper organic phase was transferred to a new glass tube. The organic solvent mix was added again for a second extraction, centrifuged, and transferred to a new glass tube. The combined 4 ml of organic phase was dried under nitrogen gas at 42°C. 150 μl of methanol was added to the dried lipids, transferred to mass spectrometry vials, and stored at -20°C until analysis by LC-MS/MS to measure the levels of the generated ceramide. The assay was linear with respect to time and protein/volume, and substrate hydrolysis was less than 10%. The reaction time and substrate concentration were controlled to ensure initial-rate conditions, enabling comparison of ASM activity across samples (**Supp. Fig. 5**).

### ACD activity assay

ACD activity assay was adapted from Bedia et al. [15]. Cells were seeded in 60-mm dishes in triplicate for each condition at a density of 1 × 10^6^ cells per dish. After 2 days, cells were harvested in 100 μl of 0.2% sucrose, pH 7.4, and sonicated for 10 s using a microtip probe sonicator at setting 2 (Heat Systems Ultrasonics, W142). The cell lysates were centrifuged at 1,500 × g for min to remove cellular debris. For the activity assay, 2 μl of RBM 14C12 substrate (Avanti Polar Lipids, Cat. No. 860855) was mixed with 73 μl of 50 mM sodium acetate buffer (pH 4.5) in each well of a black 96-well plate. Cell lysate (25 μl) was then added to each well, and the plate was incubated at 37 °C for 3 h. Following incubation, 25 μl of methanol, 100 μl of 100 μM glycine solution, and sodium periodate to a final concentration of 2.5 mg/mL were added to each well. The plate was incubated at 37 °C in the dark for an additional 1 h. Fluorescence was measured using a SpectraMax iD3 plate reader at excitation and emission wavelengths of 366 and 466 nm, respectively. Results were expressed as arbitrary fluorescence units normalized to protein content, as determined by BCA assay.

### LC-MS/MS

All samples were spiked with internal standards before extraction [16]. The same batch of internal standards was used to build the calibration curves for each species. All reported quantitation was within the linear response of the calibration curves. Samples were analyzed at the Stony Brook Biological Mass Spectrometry Core using a Thermo TSQ Quantiva mass spectrometer equipped with a Vanquish UHPLC system, and TraceFinder 5.0 analysis software. Flow: 0.5 ml/min; Mobile phase A: Fisher Water Optima LC/MS, 1 mM ammonium formate, 0.2% formic acid; Mobile phase B: Fisher Methanol Optima, 1 mM ammonium formate, 0.2% formic acid. Buffer pre-heated 30°C; Gradient: 0–1 min 80% B, 1–7.0 min 99% B, 7–16 min 99% B. The column was a Spectra 3 μm C8SR 150 × 3 mm ID HPLC Column from Peeke Scientific. Transition masses for sphingolipids, retention times, collision energies, and composition of internal standards and standards were previously published [16]. Ceramides, sphingomyelin, sphingosine, and sphingosine-1-phosphate species were reported.

### Lentivirus transfection

HEK293T cells were seeded in a 60 mm dish at 10^6^ cells/dish. The following day, 2 μg pLenti 6.3/V5 DEST with *SMPD1* [RefSeq: ID NM000543.1] or pTwist Lenti SFFV Puro WPRE and two sgRNAs mentioned below for ASM and ACD cloned in lentiCRISPRv2 were added. 1.5 µg of dVPR and 0.5 µg of VSV-G plasmid were added to 200 µl of Opti-MEM. Separately, 12 µl of Lipofectamine 2000 reagent was added to 200 µL of Opti-MEM medium. Plasmids were then added dropwise to Lipofectamine and incubated at room temperature for 20 minutes. The plasmid-Lipofectamine mixture was added to HEK293T cells dropwise to generate lentiviral particles. The medium was changed after 24 hours, and 2 days later, the lentivirus in the medium was harvested by filtering it through a 0.45 μm sterile filter. Next, 1 ml of this medium was added to 5x10^5^ MCF-7 or HeLa cells and 10 μg/ml Polybrene (Millipore Sigma) was used to increase the efficiency of viral infection. The transfectants were selected with 12 μg/ml of Blasticidin or 1μg/ml Puromycin (Invitrogen). After selection, the overexpression of the *SMPD1* mRNA and protein was confirmed by qRT-PCR, confocal microscopy, and Western blot.

### CRISPR Knockout for ASM

Two unique guide RNAs (sgRNA) were designed to target the exons in ASM using the CRISPR tool designed by Doench et al.[17], at Broad Institute. These were chosen based on the ranking provided by the sgRNA design tool. They were: (i) 5’- ACATCCCCGCACATGATGTC-3’ and (ii) 5’ -AGAGAGATGAGGCGGAGACC- 3’. The sgRNAs were cloned into lentiCRISPRv2 [18]. Furthermore, ASM knockout efficiency of the two guide RNAs was evaluated. sgRNA1 was significantly more efficient in knocking down ASM than sgRNA2, although a complete ASM knockout was not achieved (**Supp. Fig 3A**). Single-cell clones of the ASM-edited cells were obtained through Fluorescence-Assisted Cell Sorting (FACS). While several single clones showed a complete knockout of ASM mediated by sgRNA1, only one sgRNA2-edited clone showed a complete knockout (Fig 3d). The knockout percentage and the percentage of InDels in these clones were also determined using Inference of CRISPR Edits (ICE) analysis. The knockout percentage was very high (>90%) in all the clones tested (**Supp. Fig. 3B**).

ACD was knocked out using the following sgRNA sequences: (i) 5’- GAATCCATTCTAGAATACC- 3’ and (ii) 5’-CTTACCACCCTACAAAAGA-3’ *ASAH1* single- cell knockout clones were generated using the same steps as described in the paragraph above.

### Software

GraphPad Prism 10.2.3 was used to plot the graphs and for statistical tests. Inference of CRISPR Edits (ICE) analysis (https://www.editco.bio/crispr-analysis) was used to determine the InDel profile and knockout score for the two guides used to knockout *SMPD1* gene. We used the Python BioFormats and matplotlib libraries to quantify LAMP-2 imaging.

## RESULTS

### Acidic stress conditions generate PMCer

PMCer has been studied mainly for its roles in plasma membrane repair, induction of cell death, and cell adhesion. Early studies inferred PMCer generation under stress conditions using indirect methods, such as hydrolysis of plasma membrane sphingomyelin [19], and later using ceramide antibodies [20, 21]. As mentioned in the introduction, anti-ceramide antibodies, although powerful tools, provide a qualitative rather than a quantitative signal and have also been shown to recognize other lipids; their specificity is rarely validated, and a clear protocol has not been well established [22]. Some stress inducers shown to increase ceramide, presumably at the PM, include bacterial infection, chemotherapy treatment, and exposure to acidic media [23, 24]. We first screened previously reported stress conditions for their effect on PMCer levels using a method that specifically quantifies ceramide in the plasma membrane [11]. These included: Ionomycin [25], cisplatin [26], hydrogen peroxide (H_2_O_2_) [27], UV-C [28], and acidic media [29]. Confirming these previous works, all the stressors increased PMCer dramatically at the reported time and dose (**Fig. 1A**). Moreover, the increase in molar mass of PMCer was reflected in an increase in total cellular ceramide. Although the changes in total cellular ceramide were not that obvious, probably due to sphingolipid metabolism acting on distinct pools of ceramide at the same time and due to the basal PMCer constituting a small fraction of total cellular ceramide [2, 11] (**Fig. 1B** shows the loss of d18:1/16:0 and d18:1/24:1 ceramides, the most abundant species in these cells). The effect of acid pH on PMCer was much greater than any other stressor tested; therefore, it was selected to study the generation of PMCer in response to cell stress. According to the current models, PMCer originates from plasma membrane sphingomyelin hydrolysis. Of note, no statistically significant changes were observed in sphingomyelin levels under acidosis (**Supp. Fig. 1A**). However, since only a small percentage of sphingomyelin needs to be hydrolyzed to generate a large amount of ceramide, and sphingomyelin can also be generated simultaneously through de novo synthesis, these changes in sphingomyelin might be difficult to detect [1].

**Figure 1.**
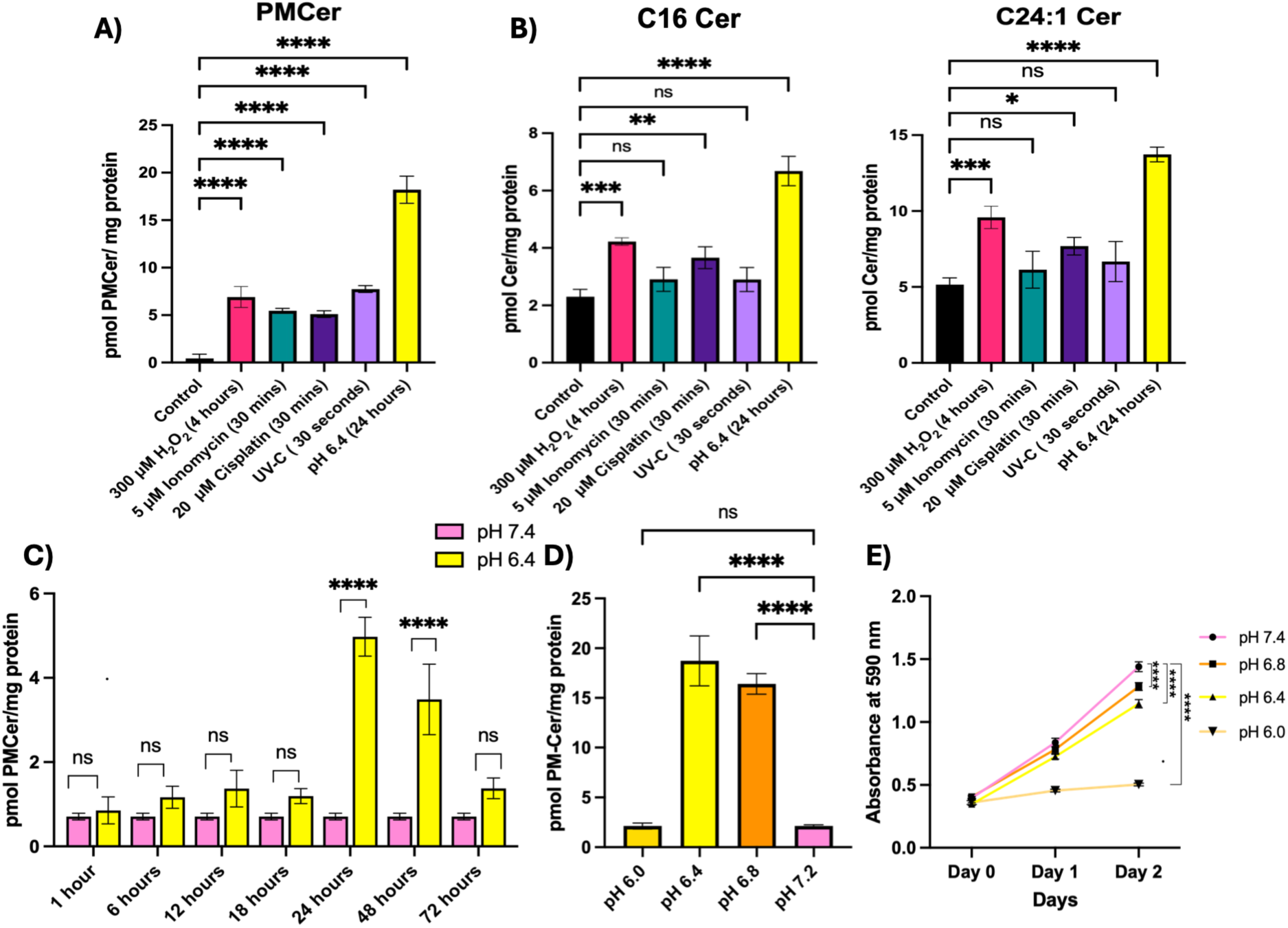
Cell stressors induced PMCer. MCF-7 cells grown in 10% FBS RPMI medium were treated with 300 µM H_2_O_2_ for 4 hours (red color), 5 µM Ionomycin for 30 mins (teal color), 20 µM Cisplatin for 30 min (indigo), and 30s of UV-C followed by 30 mins incubation (violet). Quantitative lipidomics analysis was used to calculate the mass of (**A**) PMCer and (**B**) total cellular C16 and C24:1 ceramide level. (**C**) Time course of PMCer accumulation at indicated time points under acidic stress (pH 6.4, yellow) versus neutral pH media (pH 7.4, pink), up to 72 hours. (**D**) PMCer accumulation at pH 6.0, pH 6.4, pH 6.8, and pH 7.2. (**E**) Cell viability was measured by MTT assay at the indicated pH values daily for 2 days. Statistics: 1-Way ANOVA for A, B, and D, and 2-Way ANOVA for C. ns: not significant; * p-value < 0.05; ** p-value <0.01; *** p-value < 0.001; **** p- value< 0.0001

Notably, the effects of acid on PMCer peaked at 24 hours after exposure and were maintained with a slight decrease at 48 hours, and further reduction after 72 hours (**Fig. 1C**). The optimal pH for PMCer generation was determined to be 6.4-6.8 (**Fig. 1D**). PMCer levels decreased significantly at pH 6.0, where acidic pH was affecting cell viability (**Fig. 1E**). These findings confirmed and consolidated that cellular stress induced by acid exposure stimulated the generation of PMCer.

### ASM was sufficient and necessary to generate PMCer in response to acid

We next evaluated enzymes that have previously been reported to have direct access to the plasma membrane and could be responsible for generating PMCer in response to acid. These were nSMase2 (*SMPD3*), GBA2 (*GBA2*), and SMS2 (*SGMS2),* located at the plasma membrane [19, 20], and GBA1 (*GBA1*) and ASM (*SMPD1*) [30, 31], which are reported to access the plasma membrane upon secretion into the cell media. **Fig. 2** depicts the localization of these enzymes in the cell. Of note, SMS2 is primarily involved in sphingomyelin production from Cer, but the reverse reaction has also been suggested [24].

**Figure 2.**
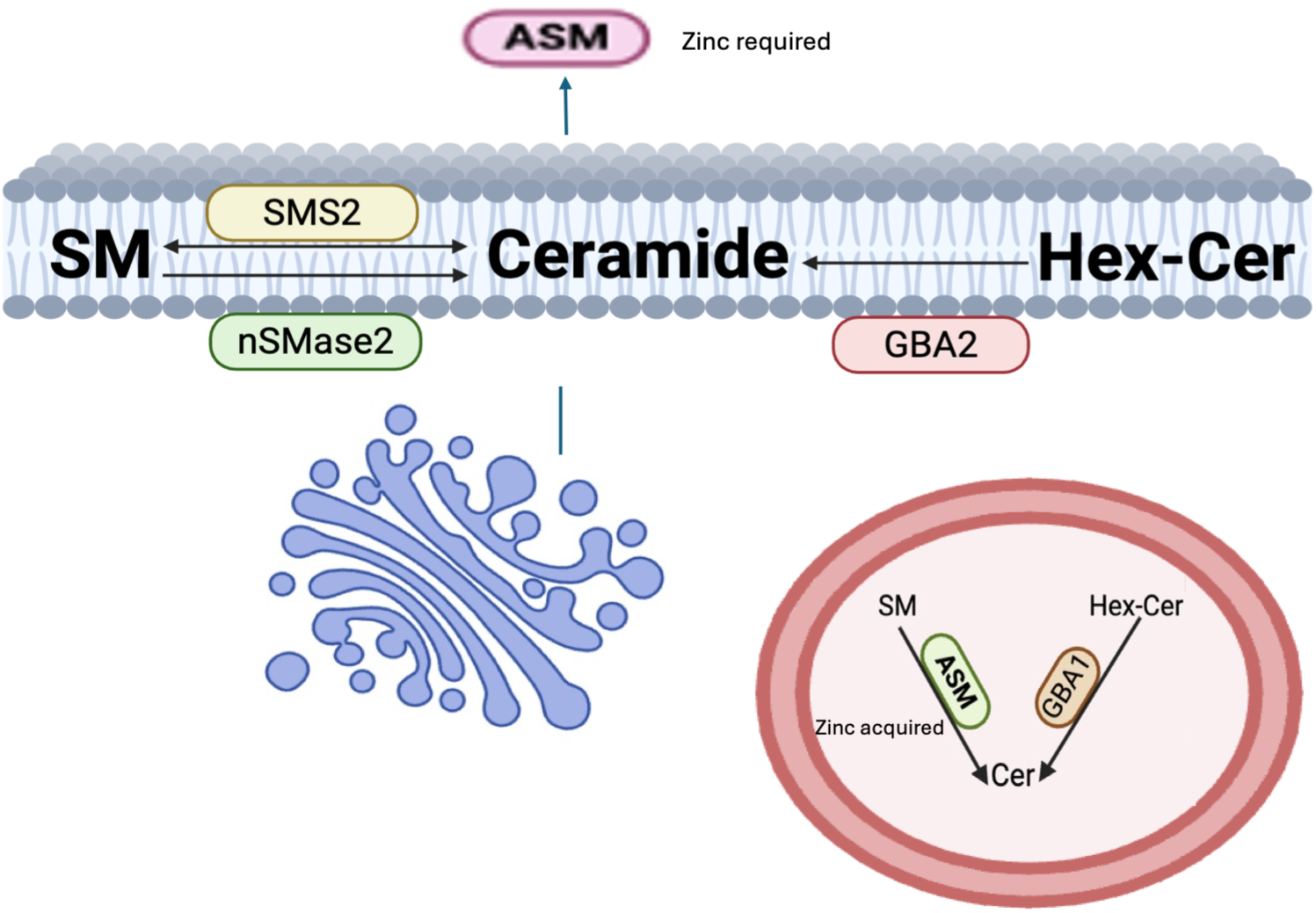
Schematic of cellular localization of nSMase2, ASM, SMS2, GBA1, and GBA2.

We recently showed that nSMase2 was the dominant enzyme regulating basal PMCer levels [7, 11], thus, we first hypothesized that nSMase2 would also be responsible for the generation of PMCer. To test this, *SMPD3* (nSMase2) was knocked down using siRNA, and consistent with published data [7], knockdown cells showed a decrease in basal PMCer. However, acidic stress did not significantly increase PMCer generation (**Fig. 3A**). Thus, we concluded that nSMase2 does not play a role in the acidic response. Therefore, to interrogate the potential role of the other enzymes, we performed siRNA-mediated knockdown of the genes in MCF-7 cells. PMCer levels in response to acid were blocked entirely upon *SMPD1* (ASM) knockdown (**Fig. 3B**). In contrast, knockdown of *GBA1*, *GBA2*, and *SGMS2* affected basal PMCer, but the cells retained the ability to generate PMCer in response to acid. These results indicated that ASM was required for PMCer generation under acidosis (confirmation of gene knockdowns encoding these enzymes is shown in **Supp. Fig. 8**).

To further confirm the siRNA screening results and the role of ASM in PMCer generation under stress, we generated CRISPR-mediated ASM knockout (KO) cells as described in the Materials and Methods. Single-cell clones of ASM KO were isolated based on the lack of ASM expression by Western blot (**Fig. 3C**). As reported by others, genetic suppression of ASM resulted in a decrease in total ceramide [32] (**Fig. 3D**), and a significant decrease in PMCer (**Fig. 3E**), confirming that ASM was required for PMCer generation under acidic conditions.

Because ASM was the only enzyme required to mediate the acid-dependent increase in PMCer, we also tested whether its overexpression was sufficient to generate PMCer. ASM overexpression through lentiviral transduction was confirmed by Western blot and immunofluorescence (**Supp. Fig. 2A and B**). As expected, overexpression of ASM increased total cellular ceramide, as represented in **Fig. 3F** for d18:1/16 and d18:1/24:1 species, although no significant changes were observed for sphingomyelin species. Overexpression of ASM exaggerated the PMCer elevation in response to acid but did not affect basal PMCer levels, suggesting ASM is not sufficient to generate basal PMCer and is only active toward PMCer generation under stress conditions (**Fig. 3G**).

### Inactive ASM mutants failed to generate PMCer

Our results so far have indicated that PMCer generation is abolished in ASM-deficient MCF-7 cells under acidic conditions. However, to definitively implicate ASM enzymatic activity is required for PMCer generation, catalytically dead ASM mutations were used to assess PMCer levels [33].

For this purpose, Niemann-Pick Disease (NPD) mutants were utilized. NPD is a lysosomal storage disorder characterized by the accumulation of sphingomyelin within lysosomes, and four types of NPD- A, B, C, and D have been identified. Of these, types A and B are caused by mutations in the *SMPD1*(ASM) gene. Several ASM mutants with varying degrees of enzymatic activity have been reported [33–36]. Among these, we selected two single-point mutants of the severe form (NPD type A)—L304P and R498L—which exhibit very low ASM activity [37].

To eliminate any residual PMCer activity from wild-type ASM, these Niemann-Pick mutants were introduced into ASM KO cells using lentiviral transduction. Single-cell clones of these mutants were established (**Fig. 4A**). Two selected clones from each of the mutants showed minimal in vitro ASM activity towards liposomal sphingomyelin (see Materials and Methods) (**Fig. 4B**). One clone for each mutant was tested for PMCer accumulation, and neither mutant restored PMCer generation at acidic pH in contrast to the action of the KO-cells reconstituted with WT ASM (**Fig. 4C**). These mutants did not increase PMCer when introduced in WT MCF-7 cells as well (**Supp. Fig 4**). Collectively, these results implicated that ASM activity was required to generate PMCer under acidic conditions.

**Figure 3.**
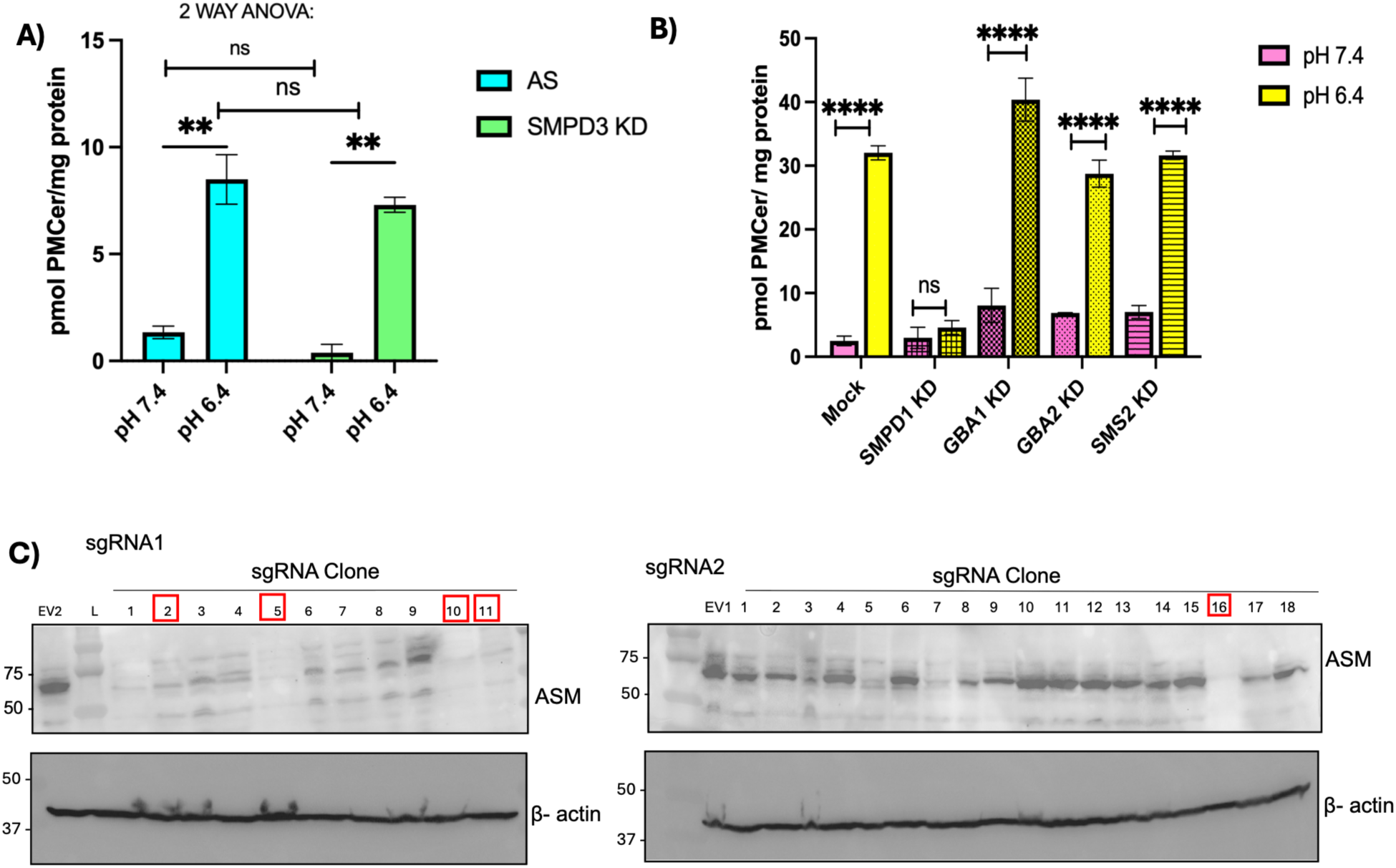
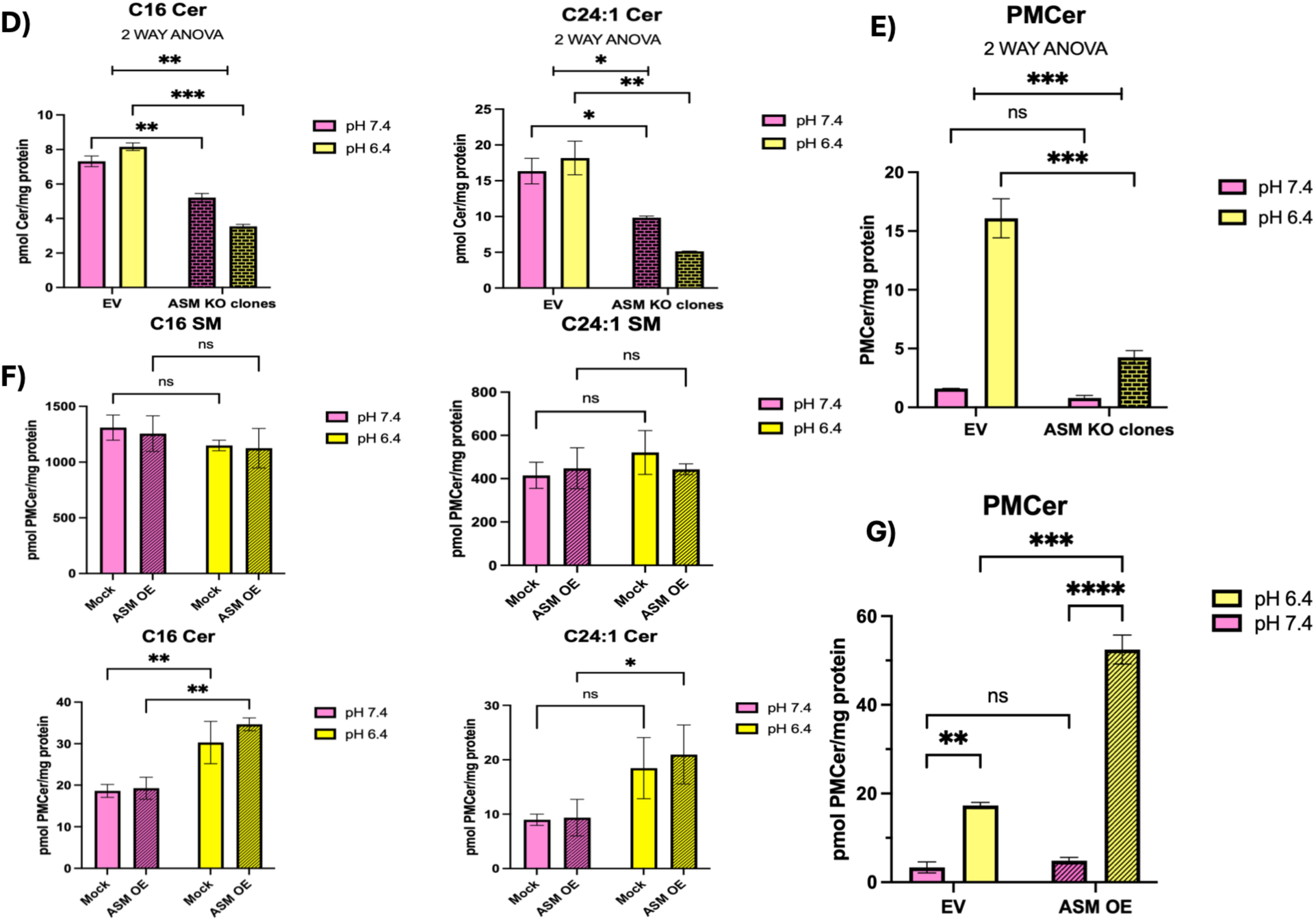
Role of PMCer metabolizing enzymes in generating PMCer under acidic stress. (**A**) PMCer levels measured in response to 24 h exposure to neutral (pH 7.4) and acidic (pH 6.4) media in control cells (AS, cyan color) and siRNA-mediated knockdown of nSMase2 (*SMPD3,* green color). (**B**) Effect of the siRNA-mediated knockdown of *SMPD1, GBA1, GBA2*, and *SGMS2* on PMCer levels at neutral pH 7.4 (pink) and acidic pH 6.4 (yellow) after 24 hours of replacing the media with the corresponding pH medium. (**C**) Single-cell clones were generated from a bulk population of ASM-knockout cells produced by lentiviral transduction. Individual cells were isolated by FACS into 96-well plates. Selected clones are indicated by red squares. (**D**) Normalized levels of C16 and C24:1 ceramides and (**E**) calculated PMCer in a pooled culture of five selected ASM (*SMPD1)* knockout clones following 24 h exposure to acidic medium (pH 6.4). The five *SMPD1* knockout clones were pooled before lipid analysis and PMCer quantification. (**F**) Levels of C16 and C24:1 cellular sphingomyelin (upper panels) and ceramide (lower panels) upon *SMPD1* overexpression after incubation for 24 hours in neutral (pH 7.4, pink) and acidic media (pH 6.4, yellow). (**G**) Effect of *SMPD1* overexpression on PMCer generation. Statistics: Two-way ANOVA. * p-value < 0.05; ** p-value <0.01; *** p-value < 0.001; **** p-value< 0.0001.

**Figure 4.**
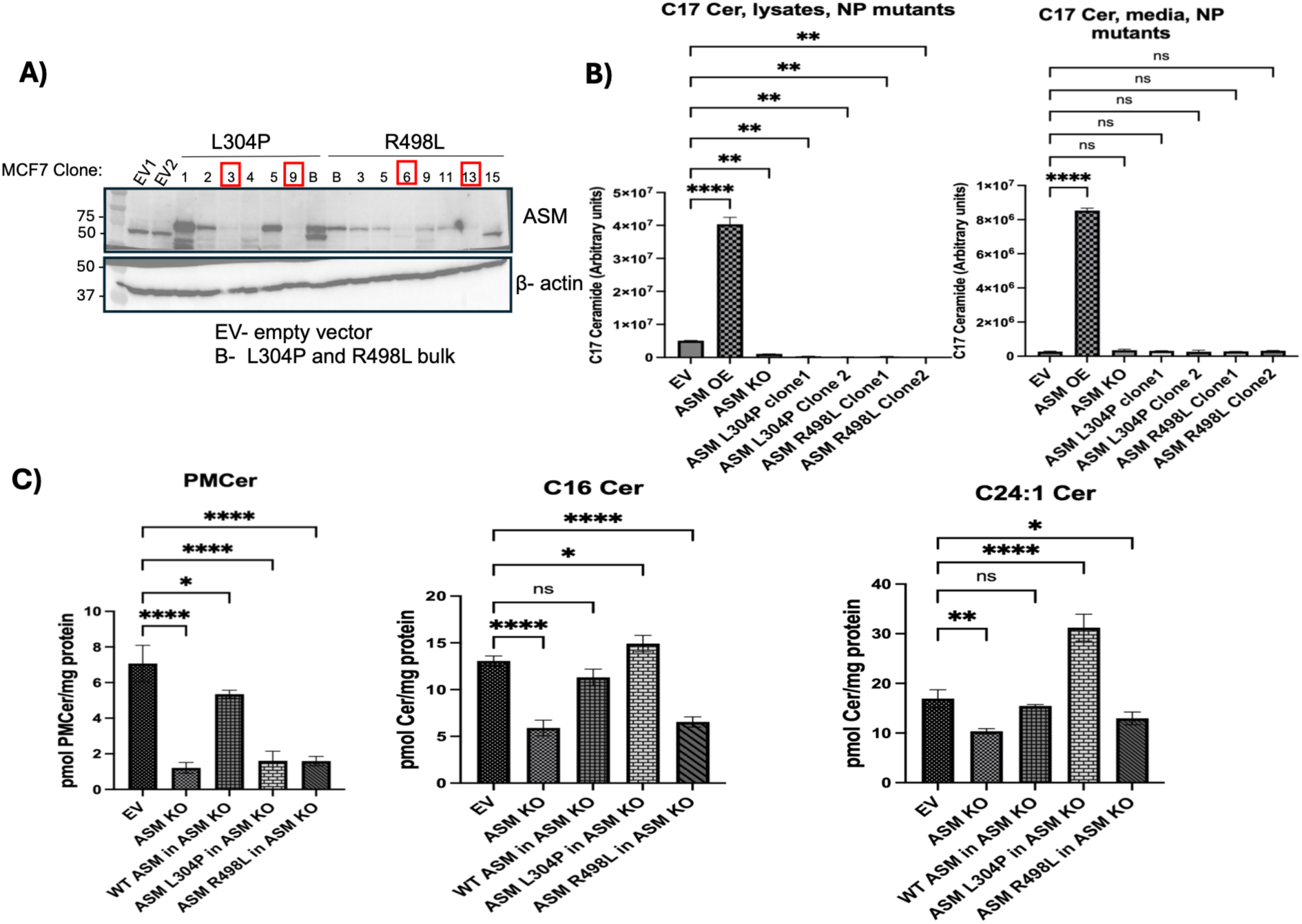
Effect of catalytically inactive single-point ASM mutants on PMCer levels. (**A**) Western blot analysis of single-cell clones expressing the Niemann-Pick (NP) disease-associated ASM mutants L304P and R498L in *SMPD1* KO cells. Single-cell clones were isolated to obtain homogeneous populations expressing each mutant. Two clones for each mutant, indicated by red squares, were selected for further analysis. (**B**) In vitro ASM activity in cell lysates (left panel) and conditioned media (right panel) from the selected clones. ASM activity was measured as C17- ceramide generation using unnatural C17-sphingomyelin as substrate. (**C**) PMCer quantification in the same L304P and R498L clones following 24 h exposure to acidic medium (pH 6.4). Empty vector (EV) was used as a control. Statistical analysis was performed by one-way ANOVA. *p- value < 0.05; **p-value < 0.01; ***p-value < 0.001; **** p-value < 0.0001.

### Enzymatically active ASM in the media does not generate plasma membrane ceramide

Two main mechanisms have been proposed for ASM regulation of PMCer in response to cell stress: Initial work showed that ASM is present either (**i**) at the plasma membrane to hydrolyze sphingomyelin to ceramide [5] or (**ii**) released into the media, which then allows access to the plasma membrane [9]. In the first mechanism, ASM was directly localized to the plasma membrane, as visualized using ASM antibodies [38]. To investigate the first mechanism, ASM subcellular localization was tracked at different time points after exposure to acidic conditions using an ASM-specific antibody, using the same protocol reported by other authors [39]. Contrary to previous works, the ASM antibody failed to show plasma membrane colocalization and showed strong intracellular co-staining with the lysosome marker lysosome-associated membrane protein 2 (LAMP2), confirming its main localization in the lysosome (**Fig. 5A**).

**Figure 5.**
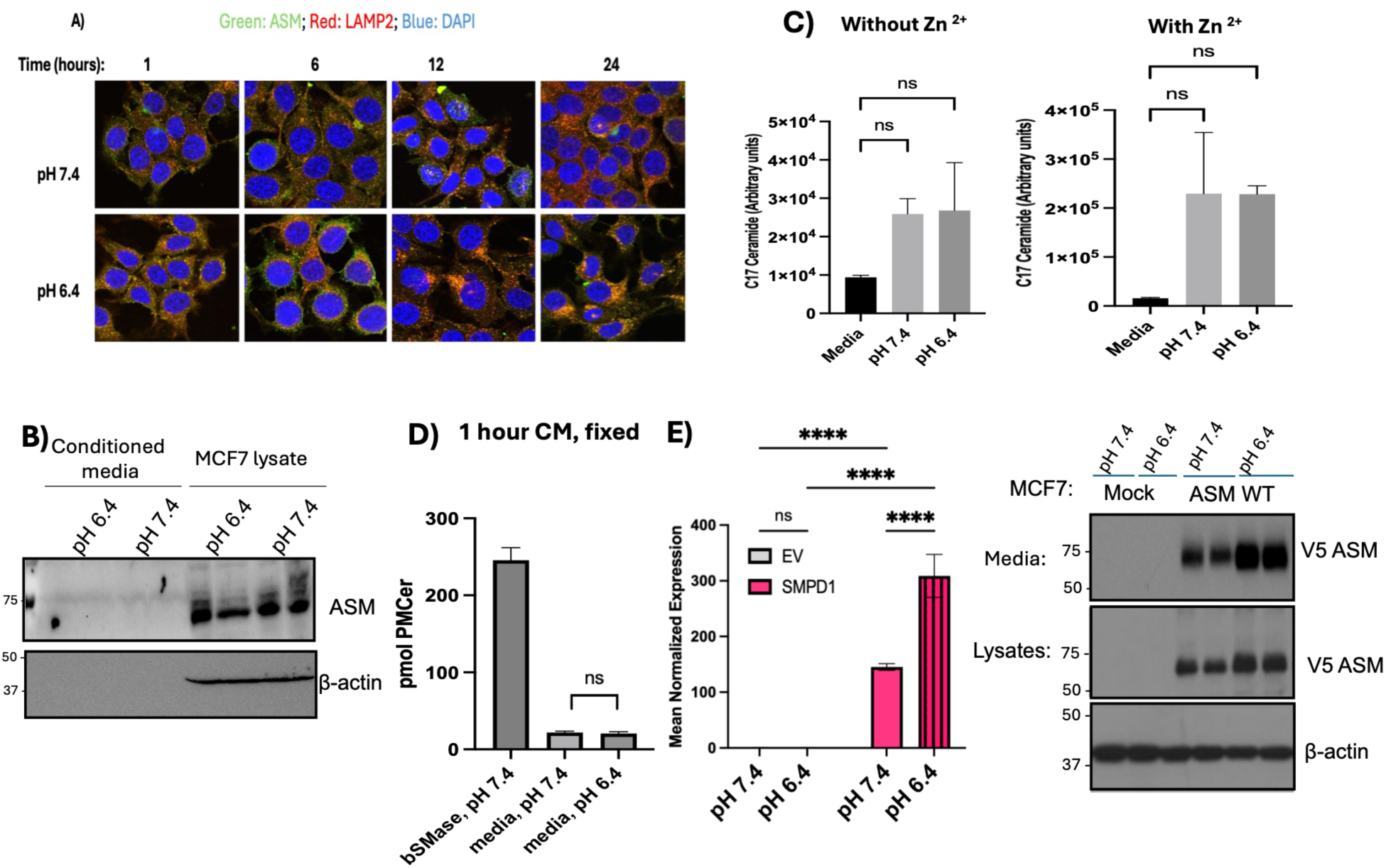
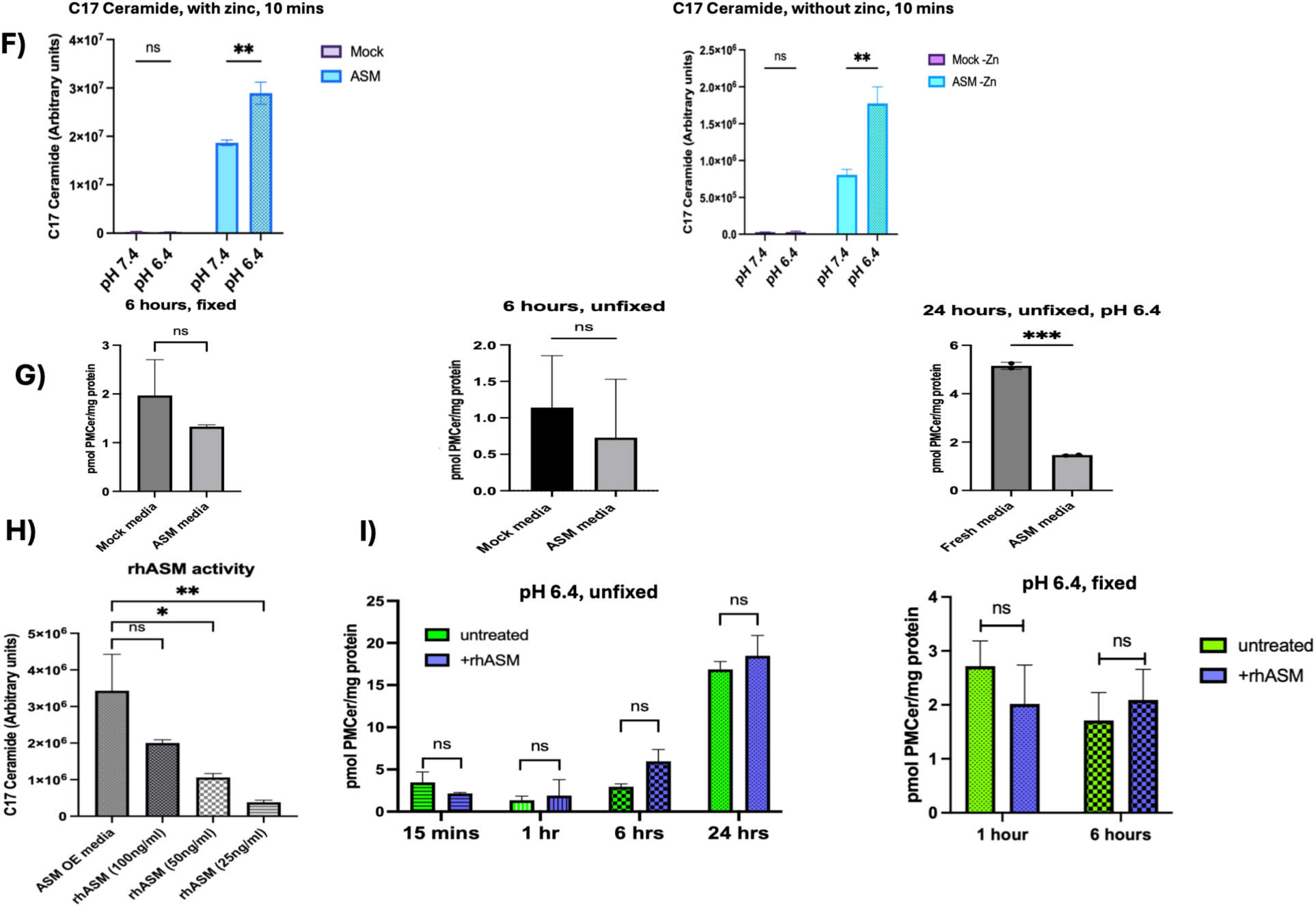
Contribution of secreted ASM to PMCer generation. **(A)** Confocal microscopy of ASM (green) and the lysosomal marker LAMP2 (red) in MCF-7 cells exposed to acidic medium (pH 6.4) for 1, 6, 12, or 24 h. Representative overlay images are shown, where yellow color represents colocalization of LAMP2 and ASM. No ASM staining was found at the plasma membrane upon acidic exposure. The nuclei of the cells are shown in blue and were stained with DAPI. **(B)** ASM protein levels in cell lysates and conditioned media from MCF-7 cells maintained in neutral media (pH 7.4) or exposed to acidic media (pH 6.4) for 24 h, assessed by Western blot using an endogenous anti-ASM antibody. Actin was used to control for equal loading in cell lysates. **(C)** ASM activity in conditioned media from MCF-7 cells maintained at neutral (pH 7.4) or exposed to acidic medium (pH 6.4) for 24 h, measured in vitro in the presence (right panel) or absence (left panel) of zinc. **(D)** Calculated PMCer levels in MCF-7 cells incubated for 24 h with conditioned media from cells maintained at neutral (pH 7.4) or exposed to acidic medium (pH 6.4) for 24 h. Recombinant bacterial sphingomyelinase (bSMase) was used as a positive control. **(E)** ASM mRNA expression (left panel) and protein levels (right panel) in ASM V5-tagged overexpressing MCF-7 cells at neutral (pH 7.4 ) and acid (pH 6.4) in cell lysates and conditioned media. **(F)** In vitro ASM activity assay of conditioned media from ASM-V5-overexpressing MCF- 7 cells (blue) and untransfected cells (Mock, purple) maintained at neutral (pH 7.4) or exposed to acidic (pH 6.4) medium for 24 h, in the presence (left panel) or absence (right panel) of zinc. **(G)** Calculated PMCer levels in MCF-7 cells incubated with conditioned media from empty vector cells (Mock) and ASM-V5-overexpressing cells for 6 or 24 h. For 6 h, cells were fixed chemically or left without fixation before being treated with conditioned media. **(H)** In vitro activity of recombinant human ASM (rhASM) at the indicated concentrations from 25 ng-100 ng rhASM/ ml. **(I)** Calculated PMCer levels in MCF-7 cells treated with rhASM for 15 min (green), 1h (gray), 6h (purple) and 24h (blue) under acidic conditions (pH 6.4). Statistical analysis: one-way ANOVA (C, G, H) and two-way ANOVA (E, F, I*). \** p-value < 0.05, **p-value< 0.01, *** p-value < 0.001, **** p-value< 0.0001.

The second proposed mechanism requires ASM to reach the extracellular space, where it could potentially hydrolyze sphingomyelin at the plasma membrane. Extracellular ASM can originate through two routes. ASM transported through the direct ER–Golgi secretory pathway is referred to as secreted ASM (S-ASM; ∼70 kDa) [31], whereas lysosomal ASM can also be released into the extracellular medium (L-ASM; ∼62 kDa) [6, 14]). In addition to their differences in molecular weight, these forms can be distinguished functionally by their requirement for exogenous zinc. Zinc is required for ASM catalysis; however, L-ASM acquires zinc during trafficking to the lysosome and therefore does not require additional zinc for activity in vitro and is referred to as the zinc-independent form. In contrast, S-ASM requires exogenous zinc for maximal activity and is commonly referred to as the zinc-dependent form [31].

To determine whether either extracellular ASM form could contribute to PMCer generation, we first examined their presence and enzymatic activity in conditioned media from MCF-7 cells under neutral and acidic conditions. Western blot analysis showed that most ASM was present in cell lysates, with substantially lower levels detected in the conditioned media (**Fig. 5B**). Exposure to acidic medium did not produce a detectable change in ASM protein abundance.

We next measured ASM activity using an in vitro assay in which liposomes containing unnatural d17-labeled sphingomyelin were used as substrate and formation of d17-ceramide was quantified as a measure of ASM activity [31]. Both zinc-dependent and zinc-independent ASM activities were detected in conditioned media. Zinc-dependent activity, corresponding predominantly to S-ASM, was approximately 10-fold higher than zinc-independent activity, corresponding to L-ASM (**Fig. 5C**). However, exposure of the cells to acidic medium did not increase the activity of either form (**Fig. 5C**). These results indicate that acidic stress does not increase the amount or activity of ASM released into the extracellular medium.

We then tested whether the ASM present in conditioned media could directly generate PMCer. Conditioned media from MCF-7 cells maintained under neutral or acidic conditions were transferred to unstimulated MCF-7 cells. Bacterial sphingomyelinase was added directly to the medium as a positive control for plasma-membrane sphingomyelin hydrolysis. Despite the presence of enzymatically active ASM in the conditioned media, neither conditioned medium induced detectable PMCer generation (**Fig. 5D**). Thus, extracellular ASM present at endogenous levels was not sufficient to hydrolyze plasma-membrane sphingomyelin and generate PMCer under these conditions.

Because these findings differed from previous reports proposing that ASM acts directly at the plasma membrane, we next tested whether a large increase in extracellular ASM could drive PMCer generation. Thus, MCF-7 cells were transduced with an ASM-expressing lentivirus, resulting in an approximately 100-fold increase in ASM mRNA. Western blot analysis confirmed marked increase in ASM protein levels in both cell lysates and conditioned media compared with control cells (**Fig. 5E**). Consistent with this increase, conditioned media from ASM-overexpressing cells displayed substantially higher ASM activity in vitro (**Fig. 5F**). However, cells overexpressing ASM showed specific activity profiles similar to those of untransfected cells (only endogenous ASM) when they were normalized to the amount of ASM mRNA expressed (**Supp. Fig. 6)**. Moreover, conditioned media from ASM-overexpressing cells showed zinc-independent increase in activity at pH 6.4 (**Fig. 5F**). This difference in activity between the neutral and acidic conditions was attributed to the higher ASM expression in acidic conditions, rather than the effect of acid itself.

We then asked whether this large increase in extracellular ASM activity was sufficient to generate PMCer. Conditioned media from ASM-overexpressing or control MCF-7 cells were applied to recipient MCF-7 cells, and PMCer generation was evaluated at 6 and 24 h. For the 6-h experiment, we examined if ASM could act directly at the plasma membrane without cellular uptake of the enzyme. To test this, the cells were chemically crosslinked with 8% PFA , as it was previously shown that crosslinking prevents internalization of exogenously applied sphingolipid metabolizing enzymes [11]. We also examined PMCer at 24-h time point because it was the earliest time at which PMCer accumulation was observed under acidic stress (**Fig. 1C**). Cells were not crosslinked at 24 h time point as it may result in lipid redistribution and confound the PMCer results.

Despite the marked increase in extracellular ASM protein and enzymatic activity, conditioned media from ASM-overexpressing cells failed to induce detectable PMCer at either 6 or 24 h by directly acting at the plasma membrane (**Fig. 5G**). Together with the results obtained using endogenous ASM levels, these findings strongly argue that extracellular ASM does not directly generate PMCer through hydrolysis of plasma membrane sphingomyelin under these conditions.

Other studies support the claim that secreted ASM generates ceramide at the plasma membrane by adding exogenous recombinant human ASM (rhASM) to culture media and reporting a decrease in total cellular sphingomyelin [40]. However, this experimental design does not prove that sphingomyelin is hydrolyzed at the plasma membrane, or that rhASM acts at the plasma membrane or by internalization into lysosomes by endocytosis. To test if enzymatically active rhASM could act directly at the plasma membrane, 100 ng/ml of rhASM was added directly into the extracellular medium following the protocol from other studies [40]. Interestingly, the in vitro activity from this amount of rhASM is in the same order of magnitude as the activity in the conditioned media from cells overexpressing ASM (**Fig. 5H**, **left panel**). MCF-7 cells were treated with rhASM in acid media from 15 min to 24 hours. Short times were chosen because ASM-dependent PMCer generation has been reported as quickly as 30 min upon cisplatin induction, while 24 h are required by acid media to increase PMCer. No increase in PMCer generation was detected at any time point after rhASM was applied to the cell media (**Fig. 5H**, **middle panel**). Of note, there was an increase in PMCer generation at 24 hours, but this effect was due to the acid treatment, and adding rhASM did not further increase PMCer. Following the same logic as before, the effect of rhASM was assessed on chemically crosslinked MCF-7 cells. The results showed that even though the rhASM was enzymatically active, it failed to generate PMCer, suggesting that ASM cannot hydrolyze sphingomyelin at the plasma membrane (**Fig. 5H**, **right panel**). Lastly, we tested if rhASM can generate PMCer at neutral pH as shown by previous studies. In contrast to these studies, we found that rhASM failed to generate PMCer at any of the tested time points at neutral pH (**Supp. Fig. 6**), providing further evidence that ASM in the media cannot generate PMCer.

### Plasma membrane ceramide is generated in the lysosomes, which fuses with the plasma membrane in response to acidic stress

Our results thus far support that ASM is necessary for PMCer generation but proved against ASM being capable of acting on the plasma membrane. Since ASM is primarily localized in the lysosome, where enzymatic conditions are favorable, one possible mechanism to account for these observations is that ASM generates ceramide in the lysosome, not at the plasma membrane, and that this lysosomal ceramide is then delivered to the plasma membrane via lysosomal exocytosis, as depicted in the model in **Fig. 6A**.

**Figure 6.**
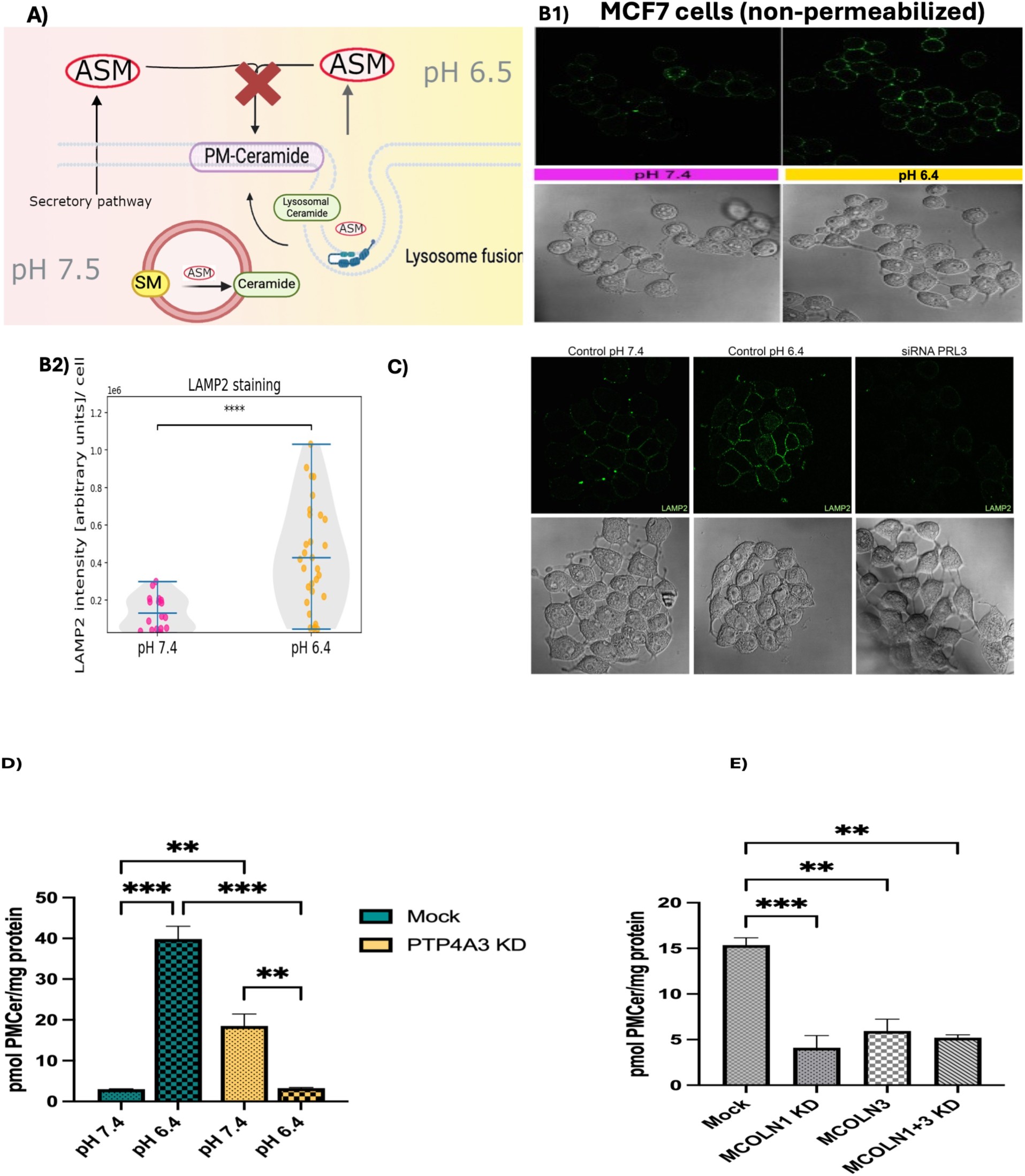
Role of lysosomal exocytosis in PMCer generation under acidic conditions. (**A**) Proposed model for PMCer generation under acidic conditions, in which lysosomal ceramide is delivered to the plasma membrane through lysosomal exocytosis. ASM in the media is not capable of hydrolyzing sphingomyelin. (**B1**) Representative confocal images showing LAMP2 (green) localization at the plasma membrane following treatment with acidic media (pH 6.4, yellow, right panel) for 24 h. Control cells are in neutral (pH7.4, pink) medium. (**B2**) Quantification of plasma membrane-associated LAMP2 following treatment with pH 6.4 media for 24 h. Quantification of LAMP2 signal/cell was calculated using the Python library BioFormats. (**C**) Effect of siRNA-mediated knockdown of *PTP4A3* on calculated PMCer levels following treatment with acidic media (pH 6.4) for 24 h. Control cells are shown in green, knockdown cells in orange. (**D**) Quantification of plasma membrane-associated LAMP2 in control and *PTP4A3*-knockdown cells following treatment with control (pH 7.4) or acidic (pH 6.4) media for 24 h. (**E**) Effect of siRNA- mediated knockdown of *MCOLN1* or *MCOLN3* on PMCer levels following treatment with acidic media (pH 6.4) for 24 h. For panels C–E, cells were transfected with the indicated siRNAs prior to acid treatment as described in the Materials and Methods. Data are presented as mean ± SEM. Statistical significance was determined by two-way ANOVA. ***p-value < 0.001; ****p-value < 0.0001.

To assess this possibility, we first determined whether the lysosomal membrane fuses with the plasma membrane in response to acidic stress. Fusion of the lysosome with the plama membrane has been monitored by tracking the presence of the lysosomal protein LAMP2 at the plasma membrane [12] [[41]. Qualitative and quantitative analysis of LAMP2 immunofluorescence showed that it was highly enriched at the plasma membrane in response to acidic media after 24 h (**Fig. 6B1 and B2)**, supporting the idea that ceramide could be transported from the lysosome to the plasma membrane. Next, to analyze the role of lysosome exocytosis on PMCer generation, the phosphatase of regenerating liver 3 protein (PRL3; *PTP4A3*) was knocked down (knockdown efficiency of *PTP4A3* is shown in **supp. Fig. 8**), as it was shown to be crucial for lysosome exocytosis specifically under acidic conditions [42]. The effect of blocking lysosomal fusion was confirmed by the absence of LAMP2 staining at the plasma membrane upon exposing the cells to acidic conditions (**Fig. 6C**). Interestingly, *PTP4A3* knockdown resulted in a significant reduction in PMCer generation only under acidic conditions (**Fig. 6D**). To further confirm the effect of lysosomal fusion on PMCer generation, the Ca^2+^ efflux channels, TRPML1/3 (encoded by genes *MCOLN1/3*), located in the lysosomal membrane, which are important mediators of lysosome exocytosis under stress, were also downregulated. Knockdown of these two channels individually or together, also reduced PMCer accumulation (**Fig. 6E**), further supporting the idea that PMCer under stress is derived from the lysosome. Knockdown efficiency of *MCOLN1* and *MCOLN3* is shown in **supp. Fig. 8**.

### Modulation of lysosomal ceramide regulates PMCer

The previous results implicated lysosome fusion was necessary to generate PMCer but also negated the need for ASM secretion into the media to generate PMCer. To demonstrate that ASM activity was required in the lysosome for PMCer generation, the ASM S508A mutant, which is reported to be active only in the lysosome and not in the media [42, 43], was used. We first confirmed that the S508A mutant was expressed in the MCF-7 cells but not secreted in the media (**Fig. 7A**). Consistent with this, the S508A mutant exhibited no activity in the media while remaining active in the cells (**Fig. 7B**). Despite its absence in the media, cells expressing the S508A mutant still generated PMCer under acidic conditions (**Fig. 7C**). This result strongly demonstrated that ASM-derived PMCer could not be generated at the plasma membrane, where the S508A mutant is not present, but ceramide had to be generated in the lysosome and transferred to the plasma membrane later.

**Figure 7.**
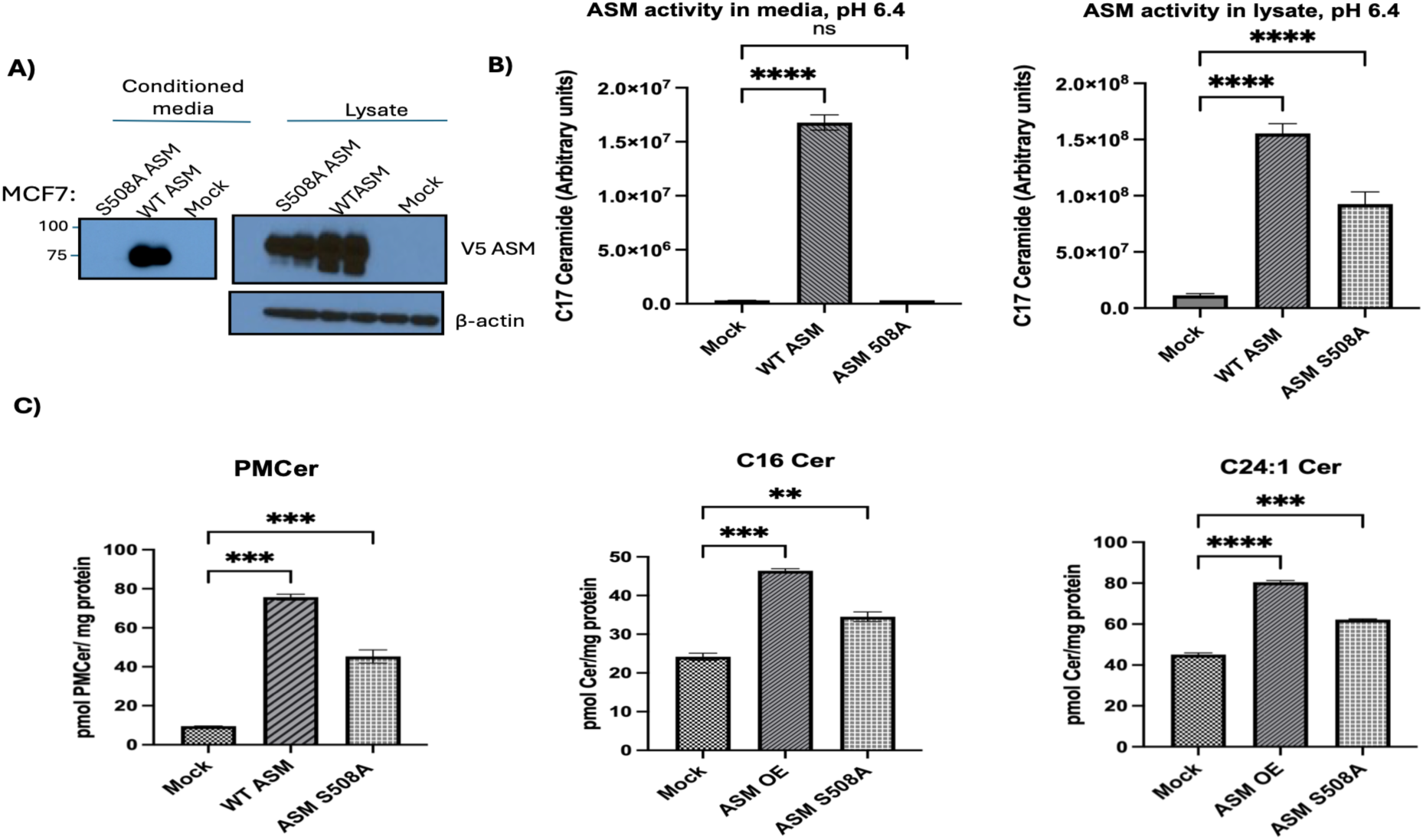
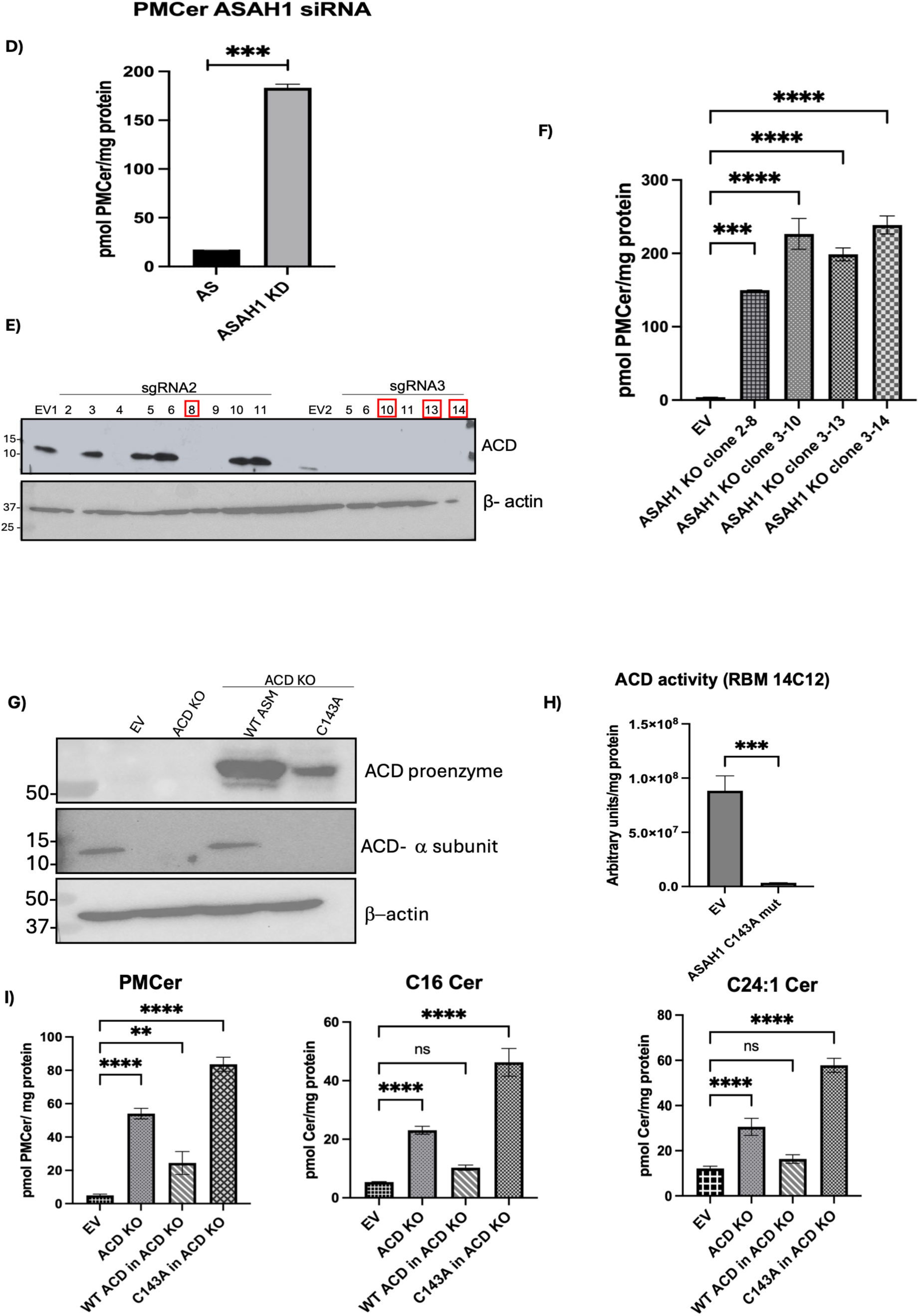
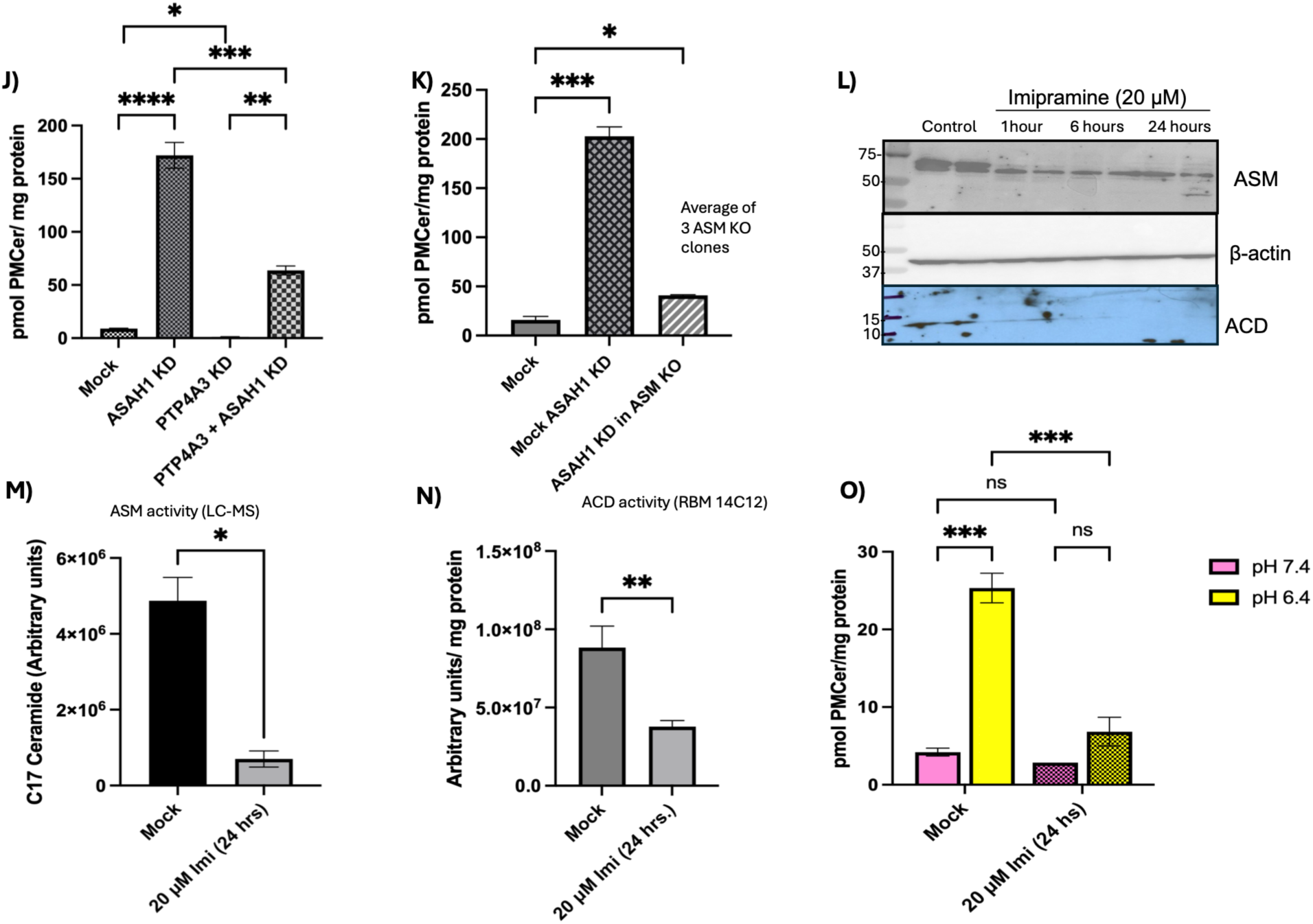
Lysosomal exocytosis contributes to PMCer generation under acidic conditions. (**A**) ASM expression levels detected by Western blotting in empty vector (Mock) and cells overexpressing wild-type (WT) ASM and the S508A mutant. The left Western blot shows conditioned media, and the right Western blot shows samples from cell lysates. (**B**) In vitro ASM activity assay from conditioned media (left plot) and cell lysates (right plot) from samples control (mock), and cells overexpressing WT and mutant S508A. (**C**) Effect of the secretion-defective ASM S508A mutant on calculated PMCer and generation of whole-cell C16 and C24:1 ceramides following treatment with acidic media (pH 6.4). (**D**) Effect of *ASAH1* knockdown on PMCer levels following treatment with acidic media (pH 6.4) (**E**) Western blot confirmation of *ASAH1* knockout in single-cell clones (**F**) PMCer levels in *ASAH1* knockout single-cell clones following treatment with acidic media (pH 6.4) (**G**) Western blot confirmation of expression of the WT *ASAH1* and catalytically inactive *ASAH1* C143A mutant introduced an *ASAH1* knockout clone by lentiviral transduction. (**H**) ACD activity in cells expressing the C143A mutant in the *ASAH1* knockout background using ceramide analog RBM14C12, as described in Materials and Methods. (**I**) PMCer levels in cells expressing the *ASAH1* C143A mutant following treatment with acidic media (pH 6.4). (**J**) Effect of combined *ASAH1* and *PTP4A3* knockdown on PMCer generation under acidic (pH 6.4) and neutral (pH 7.4) conditions. (**K**) Effect of *ASAH1* knockdown on PMCer generation in ASM knockout clones following treatment with acidic media (pH 6.4). (**L**) Time course of the effect of 20 µM imipramine treatment on ASM and ACD protein levels. (M) ASM activity following 24 h of 20 µM imipramine treatment. ASM activity was measured as described in Materials and Methods. (**N**) ACD activity following 24 h of 20 µM imipramine treatment. (**O**) PMCer generation following 24 h of 20 µM imipramine treatment. Unless otherwise indicated, cells were treated with acidic media at pH 6.4 for 24 h. Cells were transfected with 40 nM of the indicated siRNAs prior to acid treatment as described in the Materials and Methods. Statistical significance was determined by two-way ANOVA. *p-value < 0.05; **p-value < 0.01; ***p-value< 0.001; ****p-value < 0.0001.

Shifting our focus from the plasma membrane to the lysosome as the primary site of ASM activity for PMCer generation changes our understanding of how bioactive PMCer might be regulated under stress. One consequence of the proposed model, different from previous models, is that modulating lysosomal ceramide should affect PMCer levels in response to stress. It is well established that acid ceramidase (ACD; gene symbol *ASAH1*) is responsible for lysosomal ceramide catabolism, and impairment of this enzyme results in the accumulation of lysosomal ceramide, as in Farber disease [43]. Thus, depending on whether ASM acts directly at the plasma membrane or in the lysosome, ACD inhibition would produce different phenotypes. If ASM acts at the plasma membrane and generates PMCer in response to stress, *ASAH1* knockdown would not affect PMCer. However, if lysosomal ceramide is delivered to the plasma membrane, *ASAH1* knockdown would increase lysosomal ceramide and, in turn, PMCer. Remarkably, siRNA-mediated knockdown of *ASAH1* (knockdown efficiency of *ASAH1* is shown in Supp. Fig. 8) dramatically increased PMCer (**Fig. 7D**). To confirm the knockdown experiment, *ASAH1* CRISPR knockout cells (**Fig. 7E** shows selected clones) also showed a remarkable increase in PMCer levels in acidic media (**Fig. 7F**). ACD is present as the precursor form (∼50 kDa) in the cell and is cleaved to the catalytically active beta-subunit (∼35 kDa) and the smaller alpha-subunit (∼13 kDa). The beta-subunit then binds ceramide and hydrolyzes it to sphingosine. To validate that the effect of PMCer increase was due to the loss of ACD lysosomal activity, the WT ACD and a catalytically inactive C143A ACD mutant, used as a negative control, were evaluated. While the ACD precursor enzyme was expressed in both WT and C143A ACD mutant (∼50 kDa), the cleaved smaller alpha subunit (∼13 kDa) was only shown in the ASM KO cells rescued with WT ACD (**Fig. 7G**). Furthermore, the C143A mutant did not exhibit in vitro ACD enzymatic activity (**Fig. 7H**), confirming it is a catalytically inactive form. The WT ACD introduced in the knockout cells started recovering the lower PMCer phenotype, while PMCer remained high in the negative control (**Fig. 7I**). These results showed that direct modulation of lysosomal ceramide was translated to PMCer. This is consistent with our proposed model of PMCer generation and demonstrates that regulation of lysosomal ceramide is key in PMCer generation under stress and adding ACD as a non-studied player in this regulation.

We next asked whether the increased PMCer resulting from ACD loss still required lysosomal exocytosis. ACD depletion was therefore combined with depletion of PRL3 (*PTP4A3*), which inhibits lysosome fusion with the plasma membrane. The increase in PMCer caused by ACD loss was significantly reduced following *PTP4A3* depletion (**Fig. 7J**), indicating that accumulation of lysosomal ceramide alone is insufficient and that lysosomal exocytosis is required for its appearance at the plasma membrane. Furthermore, depletion of ASM in ACD-knockout cells abolished the PMCer accumulation caused by ACD loss (**Fig. 7K**). Together, these results place ASM and ACD upstream of lysosomal exocytosis and support a model in which the lysosomal ceramide pool is generated by ASM, regulated by ACD, and subsequently delivered to the plasma membrane.

As an independent approach, we used imipramine, which reduces lysosomal ASM and ACD activity by disrupting their association with lysosomal membranes and promoting their proteolytic degradation by cathepsins [24, 44][45]. Imipramine treatment reduced ASM protein levels within 1 h, whereas ACD protein was strongly reduced by 6 h, with suppression of both enzymes persisting for up to 24 h (**Fig. 7L**). Correspondingly, cellular ASM and ACD enzymatic activities were significantly reduced (**Fig. 7M,N**), consistent with previous reports [46]. Importantly, imipramine treatment abolished PMCer accumulation under acidic conditions (**Fig. 7O**). These findings provide an independent pharmacological confirmation that lysosomal sphingolipid metabolism is required for stress-induced PMCer generation.

Together, these complementary approaches support a model in which extracellular ASM is not required for PMCer generation. Instead, ASM generates ceramide within the lysosomal compartment, where its abundance is counter-regulated by ACD. In response to acidic stress, lysosomal exocytosis delivers this ceramide-enriched membrane to the plasma membrane, resulting in the acute accumulation of PMCer.

## Discussion

ASM has been implicated in cellular stress responses through a model in which it translocates to the plasma membrane and by acting on SM in the outer leaflet of the plasma membrane generates PMCer, a bioactive lipid involved in stress signaling. However, extracellular conditions are not optimal for ASM activity, and direct ASM-mediated hydrolysis of sphingomyelin at the plasma membrane has been assumed rather than demonstrated. We propose an alternative mechanism in which lysosomal fusion with the plasma membrane increases PMCer by transferring lysosome-derived ceramide to the plasma membrane. This model is consistent with previous observations showing that ASM is required for stress-induced increases in PMCer, but the model proposed here does not require ASM activation, secretion, translocation, or catalytic activity at the plasma membrane. Thus, ASM contributes to PMCer generation indirectly, through lysosomal ceramide production and subsequent membrane fusion, rather than by acting directly at the cell surface.

Consistent with the recurrent implication of ASM in cellular stress responses, we identified extracellular acidification as a stress condition that induces PMCer accumulation in an ASM-dependent manner. We therefore used acid stress as a defined model to interrogate the mechanism by which ASM regulates PMCer during cellular stress. Among other enzymes implicated in PMCer metabolism, modulation of nSMase2, GBA2, or SMS2 did not reproduce the ASM-dependent response to acid stress. This distinction is particularly relevant for nSMase2, which we recently showed regulates steady-state PMCer under physiological conditions, including confluence, extracellular vesicle formation, and cytokine signaling [47, 48], but has not been clearly implicated in stress-induced PMCer generation [7, 49, 50]. For GBA2 and SMS2, their proposed roles in PMCer regulation remain largely hypothetical, and their contribution to PMCer-dependent biological responses is not well established. Thus, extracellular acidification provides a tractable system to specifically investigate how ASM contributes to stress-induced PMCer accumulation.

A central question is how the prevailing model of ASM-mediated PMCer generation can be reconciled with the biochemical constraints on ASM activity outside the lysosome. Early studies detected ASM at the plasma membrane in response to stress using antibody-based imaging but did not establish how ASM reached or associated with the cell surface, or whether membrane-associated ASM was active in this new compartment. Subsequent studies proposed secretion of ASM, and later lysosome–plasma membrane fusion, as mechanisms by which ASM could reach the extracellular surface. Because these events were accompanied by increased PMCer, it was inferred that extracellular ASM hydrolyzed sphingomyelin in the outer leaflet of the plasma membrane. However, localization or secretion of ASM does not demonstrate catalytic activity at this site, and several biochemical properties of the enzyme make such activity difficult to reconcile with extracellular conditions.

ASM is optimized for the lysosomal environment, where acidic pH favors the protonation states required for catalysis and promotes interaction with anionic lipids such as bis(monoacylglycero)phosphate (BMP) [51–53]. In contrast, the outer leaflet of the plasma membrane is enriched in neutral lipids such as sphingomyelin and phosphatidylcholine and is exposed to a near-neutral, relatively high-ionic-strength extracellular environment. These conditions are unfavorable for ASM membrane association and catalytic activity. Phosphatidylserine (PS) exposure during apoptosis has been proposed to provide an anionic surface for ASM binding; however, ASM interaction with PS and other anionic lipids has primarily been demonstrated in model membranes under acidic conditions rather than at the surface of intact cells [45]. Secretory ASM also has a greater dependence on Zn²⁺, further constraining its activity extracellularly. Although secretory ASM can hydrolyze oxidized sphingomyelin in low-density lipoproteins at neutral pH, this activity is substantially lower than under acidic conditions and it depends on the more accessible SM exposure[54]. in support of these biochemical constraints, our results show that recombinant human ASM and ASM from conditioned medium hydrolyzed sphingomyelin-containing liposomes under favorable conditions but failed to generate PMCer at the surface of intact cells.

Our model resolves these apparent contradictions without requiring ASM to function under unfavorable extracellular conditions. Under acid stress, our data demonstrate that ASM generates ceramide within the lysosomal compartment and that lysosome–plasma membrane fusion subsequently transfers this ceramide to the plasma membrane. Thus, lysosomal fusion, rather than activation or translocation of ASM at the cell surface, emerges as the regulated step controlling PMCer accumulation. In this model, ASM remains essential because it determines the lysosomal ceramide available for transfer, while alterations in ASM, ACD, or other regulators of the lysosomal ceramide pool can modulate the magnitude of PMCer accumulation. Accordingly, mechanisms controlling lysosome–plasma membrane fusion become key regulators of stress-induced PMCer.

Importantly, ACD provides reciprocal control over this pathway. ACD knockdown produced a marked increase in PMCer, indicating that degradation of lysosomal ceramide limits the amount ultimately delivered to the plasma membrane. The requirement for ACD catalytic activity was further supported using the catalytically inactive C143A mutant in *ASAH1*-deficient cells. Thus, PMCer accumulation reflects not only lysosomal fusion but also the balance between lysosomal ceramide generation and degradation. This point may be particularly relevant in pathological settings in which ASM or ACD activity is altered, including lysosomal storage disorders and cancer. More broadly, our data shift the focus from ASM activity at the plasma membrane to regulation of the lysosomal ceramide pool and lysosome–plasma membrane fusion as determinants of acute PMCer signaling.

In summary, we identify a pathway in which lysosome-derived ceramide is transferred to the plasma membrane during stress-induced lysosomal exocytosis, generating an acute increase in PMCer. This mechanism provides an alternative explanation for the long-standing association between ASM and stress-induced PMCer while avoiding the requirement for ASM catalytic activity under unfavorable extracellular conditions. More broadly, it establishes regulated organelle-to-plasma-membrane lipid transfer as a mechanism for rapidly modifying the abundance of a bioactive lipid at the cell surface.

## Acknowledgements

The authors wish to acknowledge the Stony Brook Cancer Center Biological Mass Spectrometry Shared Resource for expert assistance with Lipidomics analysis. We would also like to thank Dr. Chiara Luberto for suggesting important experiments to improve the study. We would also like to thank the members of the Lipid Cancer Laboratory for their input on the project.

## Author contributions

D. C. and A.K. writing–original draft; D. C. supervision; D. C. resources; D. C. and Y. A. H. funding acquisition; D. C. and A.K. formal analysis; D. C. and A.K. data curation; D. C., Y. A. H., M. D., and A.K. conceptualization; D. C., A.K., A. G. O., M. J. H., and R.C. investigation.

## Funding information

DC: Carol M. Baldwin Foundation for Breast Cancer Research, Cancer Center at Stony Brook Start-Up, and National Cancer Institute (NCI), Grant/Award Number: CA097132. YAH: National Cancer Institute (NCI), Grant/Award Number: CA218678 and CA097132.

## Notes

### Competing Interest Statement

The authors have declared no competing interest.

