## Supplemental figures for "Acid Stress-induced plasma membrane ceramide arises from lysosome-plasma membrane fusion"

Supp Fig 1:

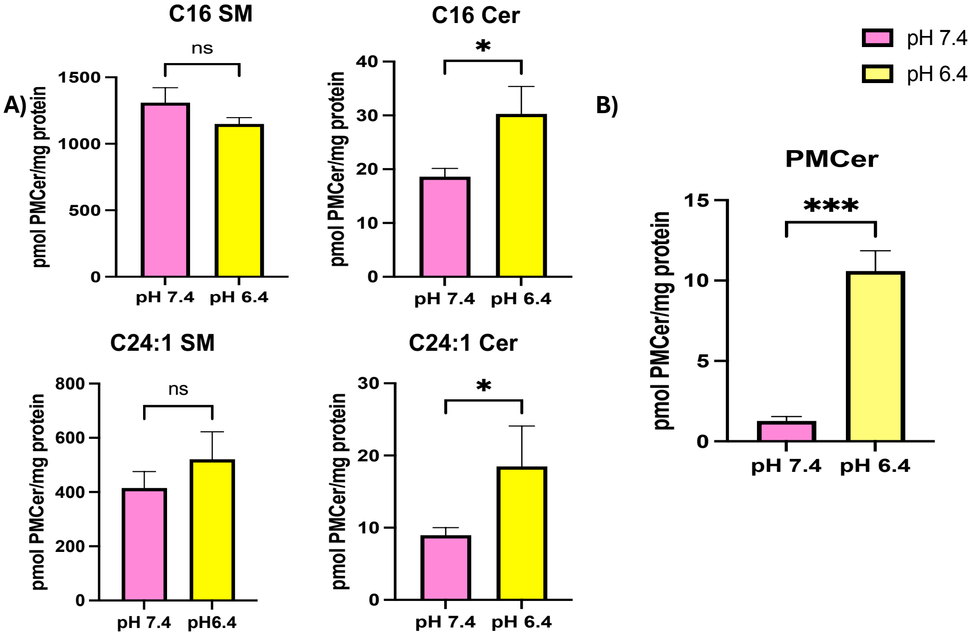


**Supplementary Figure 1. Changes in sphingomyelin, ceramide, and PMCer levels under acidic conditions.**

(A) Levels of C16 and C24:1 sphingomyelin and ceramide following treatment with acidic media (pH 6.4) for 24 h. (B) PMCer levels following treatment with acidic media (pH 6.4) for 24 h. Statistical significance was determined by two-way ANOVA. ns, not significant; * p-value < 0.05; **p-value< 0.01; ***p-value < 0.001; ****p-value < 0.0001.

Supp. Fig. 2

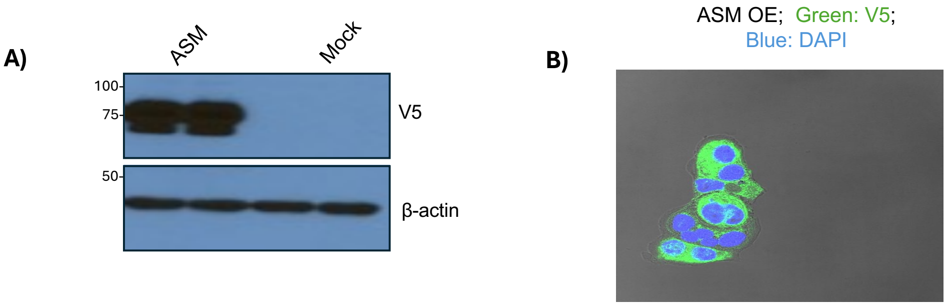


**Supplementary Figure 2. Overexpression and localization of ASM in MCF-7 cells.**

ASM protein overexpression was confirmed by immunoblotting. (B) Confocal microscopy showing the subcellular localization o

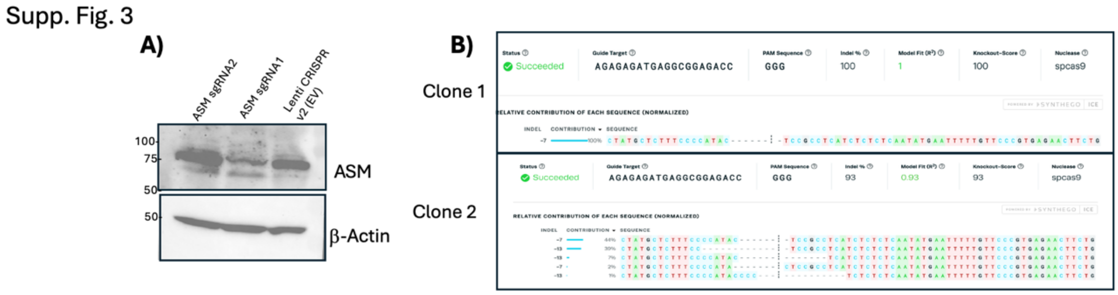


f overexpressed ASM (green).

**Supplementary Figure 3. Confirmation of ASM knockout in MCF-7 cells.**

Confirmation of ASM (*SMPD1*) knockout by immunoblotting in the bulk MCF-7 population. **(B)** ICE analysis of two representative *SMPD1* knockout clones generated using sgRNA1 and sgRNA2, respectively, showing the knockout efficiency and percentage distribution of individual InDels in each clone.

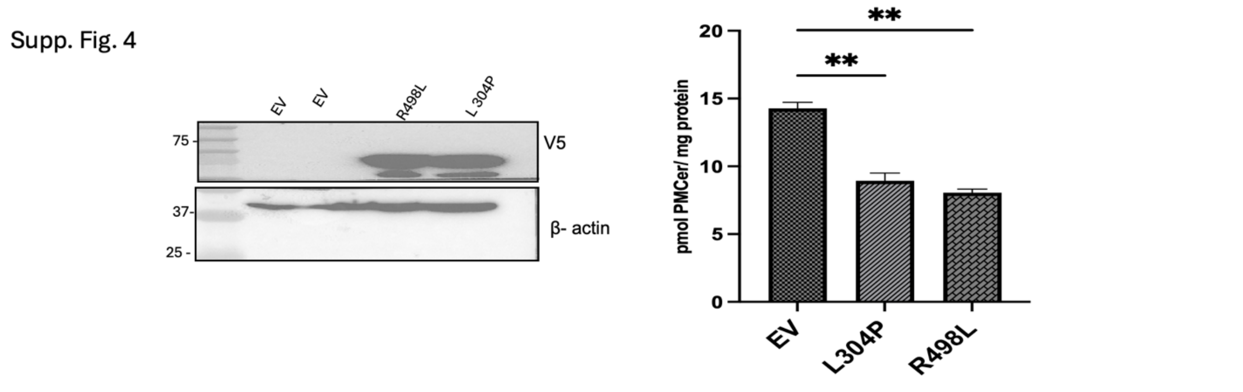


**Supplementary Figure 4. Niemann–Pick disease-associated ASM mutants do not increase PMCer generation.**

MCF-7 cells expressing wild-type ASM or the Niemann–Pick disease-associated ASM mutants L304P or R498L were analyzed for PMCer generation under acidic conditions (pH 6.4) for 24 hours. Expression of either mutant did not increase PMCer generation compared with the empty-vector (EV) control.

Supp. Fig. 5

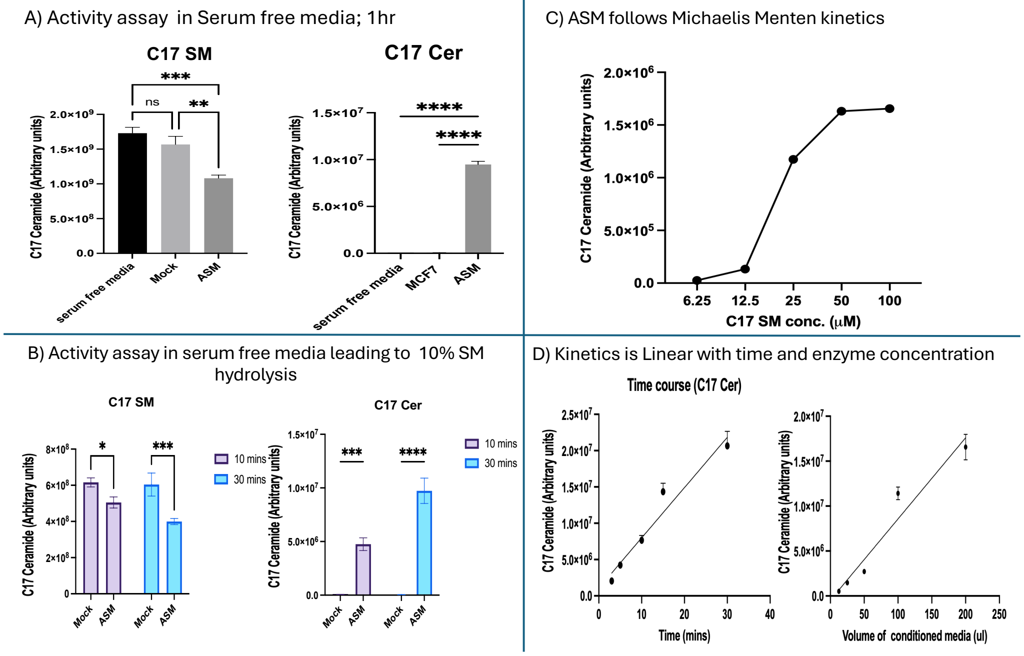


**Supplementary Figure 5. Optimization and characterization of the ASM activity assay in conditioned media.**

ASM activity measured in serum-free conditioned media. (B) Optimization of the incubation time for the ASM activity assay. (C) Enzyme kinetics of ASM activity measured in conditioned media. (D) Effect of assay time and conditioned-media volume on measured ASM activity. Because the exact concentration of ASM in conditioned media could not be determined, conditioned-media volume was used as a proxy for enzyme input. Statistical significance was determined using one-way or two-way ANOVA, as appropriate. ns, not significant; *p-value< 0.05; **p-value < 0.01; ***p-value < 0.001; ****p-value < 0.0001.

Supp. Fig. 6

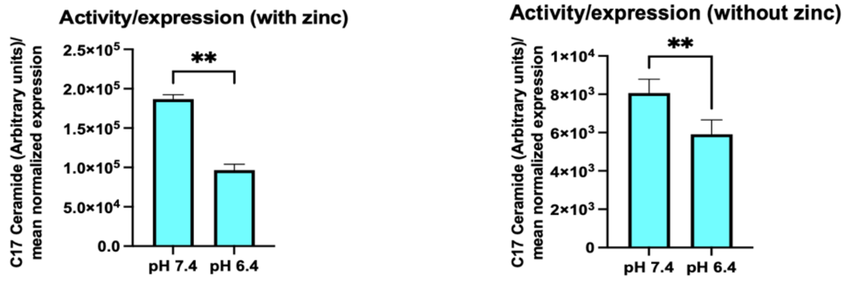


**Supplementary Figure 6. Effect of zinc on the activity of recombinant ASM stably overexpressed in MCF-7 cells.**

ASM activity in MCF-7 cells stably overexpressing recombinant ASM was measured in the presence or absence of 0.1 mM zinc and normalized to ASM expression as described in Materials and Methods.

Supp. Fig. 7

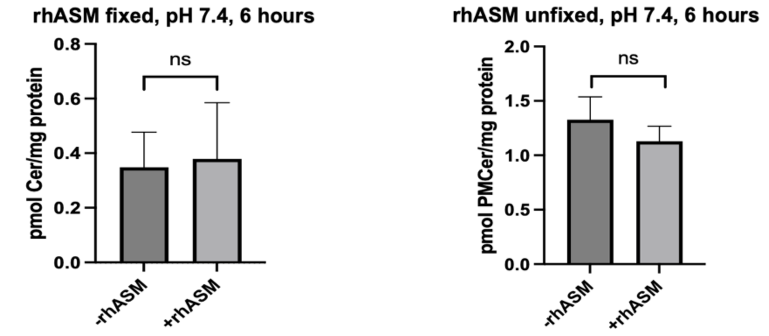


**Supplementary Figure 7. Effect of recombinant human ASM on PMCer generation under neutral conditions.**

Effect of 100 ng/ml recombinant human ASM (rhASM) treatment on PMCer generation under neutral conditions (pH 7.4) following treatment for 6 h and 24 h.

Supp. Fig. 8

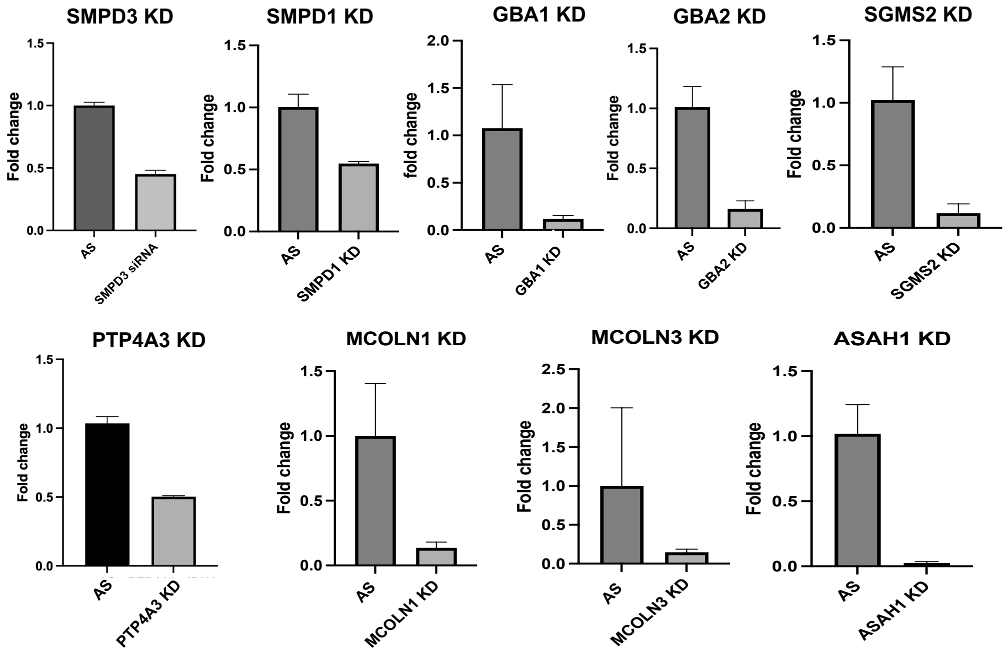


**Supplementary Figure 8. Knockdown efficiencies of genes encoding sphingolipid metabolizing enzymes and proteins involved in lysosome exocytosis machinery.**

mRNA levels of *SMPD1*, *SGMS2, GBA1*, *GBA2, PTP4A3, MCOLN1, MCOLN3* and *ASAH1* following siRNA-mediated knockdown, measured by RT-qPCR. Cells were transfected with the indicated siRNAs prior to acid treatment as described in the Materials and Methods.
